# Reversible DNA Translocation as a molecular caliper to probe the nanoscale asymmetry of glass nanopores

**DOI:** 10.1101/2025.07.11.664396

**Authors:** Sukanya Sadhu, Gautam V. Soni

## Abstract

Studying DNA conformation is crucial for understanding gene regulation, chromatin structure, and genomic stability. Nanopores have proven to be excellent label-free, high-throughput tools for studying conformational changes in various biomolecular structures. Conical glass nanopores are widely used in solid-state nanopore studies due to their simple and cost-effective fabrication. However, their inherent geometric asymmetry introduces distinct characteristics in the translocation dynamics of DNA. Here, we demonstrate bi-directional translocation of multiple DNA lengths through a conical nanopore to understand the role of pore asymmetry. We show a quantitative comparison of various parameters, such as the conductance drop (ΔG), translocation time (Δt), event charge deficit (ECD) and percentage of DNA folding in both directions. A natural output of our ECD-based analysis is the estimation of the effective sensing length of our conical pore geometry. In the nanoscale regime, sensing length controls spatial resolution of the nanopore detector and is challenging to measure. Our study reveals significant experimental insights into the dependence of DNA length, translocation directionality, and applied voltage on the translocation mechanism, contributing to a broader utilization of glass nanopores in sensing technologies.

**TOC Figure:** 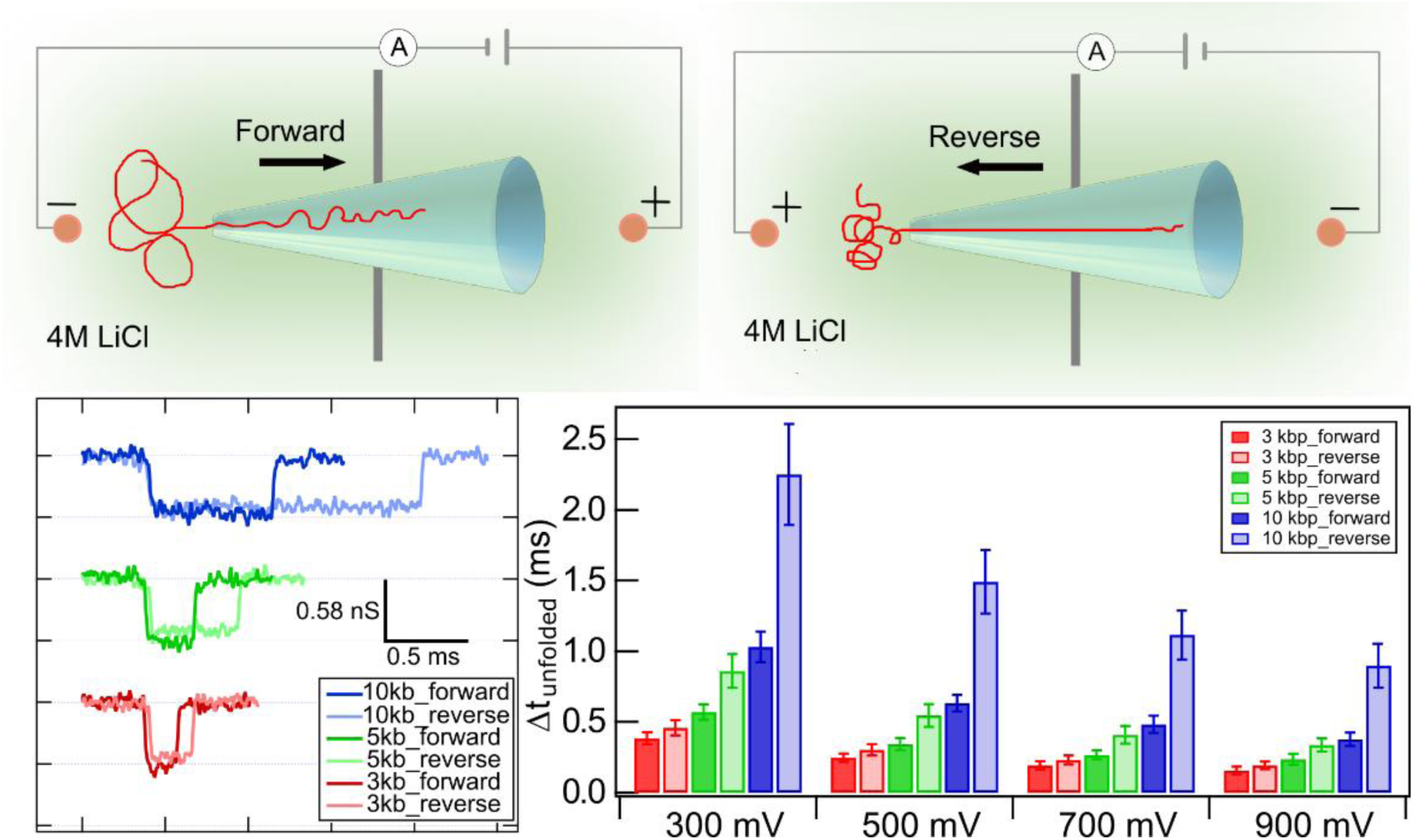

## INTRODUCTION

Studying the transport of biopolymers such as DNA through a nanoscale confinement is crucial for revealing how DNA interacts with confined environments^1^, mimicking conditions in cellular pores and channels^2,3^. This aids in understanding DNA-protein interactions^4–7^, molecular transport mechanisms^8^, and the influence of confinement on biological processes^9^ like gene regulation and chromatin organization^10^. Nanopores are particularly useful for this purpose due to their high throughput and ability to resolve distinct molecular conformations^11–13^. The basic principle of nanopore functioning is based on the resistive pulse technique^14,15^. Here, the nanopore constitutes the ionic path between the two fluid chambers filled with salt solution. On application of a voltage bias across the nanopore, ion current flows through the open nanopore. When an analyte molecule, e.g. DNA or protein, approaches the pore, it is electrophoretically driven through the pore. During translocation through the nanopore, the blockage of the ionic current proportional to its own volume is measured as its electrical signature. By investigating the blockage amplitude and duration, the size, length, and charge estimation of the molecule can be done. Thus, nanopore offers an effective label-free platform for probing the physical behaviour of DNA translocating through confinement^1,16,17^.

Nanopores are broadly categorized into biological and solid-state types, each with distinct fabrication processes and application scopes. Biological nanopores are formed from natural proteins and are characterized by precise structures and interfacial chemistries, offering high specificity for certain molecules^18–21^. However, they suffer from limitations such as low mechanical strength, fixed pore sizes and strict environmental requirements, which can restrict their practical applications^22^. Solid-state nanopores, which include both planar and conical glass nanopores, are fabricated using techniques such as focused ion beam milling, controlled dielectric breakdown^23–25^ or laser-assisted pulling of glass capillaries^26^. Solid-state nanopores exhibit uniformity and reproducibility, making them suitable for high-throughput screening^27^. Planar nanopores may generate higher noise levels due to their flat geometry^22^. Conversely, conical glass nanopores, known for their tapered structure, demonstrate enhanced sensitivity and lower noise due to better ionic flow characteristics^28–30^. While conical nanopores provide superior detection capabilities with enhanced rectification properties, their fabrication and characterization can be more complex^31^. The inherent asymmetry of conical nanopores introduces distinct behaviours in bi-directional DNA transport^32–38^. Most studies with conical glass nanopores have predominantly focused on forward translocation, where the DNA molecules are driven from the bulk solution into the conical nanopore under the applied voltage. In contrast, reverse translocation, wherein DNA inside the capillary exits the confinement into the bulk solution, remains relatively under-explored despite its unique advantages. Previous studies^34^ have shown the possibility that in reverse translocation, DNA moves at a slower speed, which may enhance temporal resolution. Additionally, reverse translocation reduces the occurrence of DNA folding, which is particularly beneficial for studying protein-DNA complexes that are sensitive to folding artefacts. A comprehensive study of various factors such as nanopore geometry, external electric fields, and surface interactions that affect DNA translocation dynamics remains incomplete. Key parameters like conductance change (ΔG), translocation time (Δt), DNA folding, and event charge deficit (ECD) are influenced by these factors, yet their directional dependencies and interplay with nanopore asymmetry are not fully characterized. To address these knowledge gaps, it is critical to systematically study and compare forward and reverse translocation processes, particularly in asymmetric nanopores like conical glass structures.

An important concept in nanopore sensing is the ‘effective sensing length’ (L_eff_), defined as the distance along the nanopore axis where the drop in ionic conductance due to a passing analyte can be detected over the nanopore noise^39^. The voltage applied across the nanopore develops a strong electric field region inside the pore, dropping sharply outside the mouth of the pore and more gradually along the conical length of the nanopore. This high field region, where the field drops to 1/e of its maximum strength on either side of the nanopore, is termed as the effective sensing length^40^. The sensing length governs the spatial resolution of a nanopore. In planar solid-state and biological nanopores, the sensing region is well defined by the membrane thickness and protein channel length, respectively^41,42^. In contrast, the sensing region in conical glass nanopores extends deeper into the pore body and is less well-defined. Factors such as fabrication variability, surface roughness, diameter, and cone angles all contribute to heterogeneity in pore geometry and hence sensing region^43,44^. A direct measurement of L_eff_ via SEM visualization of the internal geometry of the conical nanopore remains elusive^14,43,45^, making its *in situ* measurement important for the progress of this field.

In our present work, we aim to bridge this gap by systematically investigating the bidirectional translocation of linear DNA through conical glass nanopores. We prepared and used linear DNA of varying lengths (3, 5 & 10 kbp) and conducted experiments at multiple forward and reverse voltage biases (± 300 mV, ± 500 mV, ± 700 mV, ± 900 mV). We have done a quantitative analysis and comparison of various parameters such as conductance drop (ΔG), translocation time (Δt), event charge deficit (ECD), and percentage of folding in both directions. Our data elucidates distinct differences in these parameters between the two translocation directions. Using ECD-based analysis, we determined the fold lengths of individual DNA strands and then calculated L_eff_ from them. Additionally, we employed a double-cone analytical model^43^ to independently calculate the sensing length and found strong agreement between our experimental and theoretical results. Interestingly, our findings also reveal notable trends in L_eff_ as a function of voltage, suggesting that the sensing region expands with increasing electric field strength—a phenomenon with important implications for nanopore-based sensing applications. A key experimental challenge in solid-state nanopore studies is the significant pore-to-pore variability, which makes quantitative comparison difficult. To minimize this variability, we have used the same nanopore across a systematic series of experiments, varying the applied voltage, DNA length, and translocation direction within the same device. This controlled approach reduces variability by isolating the influence of each parameter, enhancing data reliability, and enabling a more precise comparison and interpretation. Furthermore, to ensure robustness, these experiments were repeated across different nanopores to further validate the data trends. Our findings enable a comprehensive characterization of DNA translocation parameters through nanoscale confinement, along with precise measurement of nanopore sensing length, with applications in high-resolution biomolecular sensing and future nanopore-based sensor technologies.

## MATERIALS AND METHODS

### Linear DNA Preparation

In our nanopore translocation experiments, linear DNA of lengths 3 kbp, 5 kbp, and 10 kbp was used. All three plasmids were amplified by growing in E. coli DH5α overnight on 180 rpm shaker at 37 °C and extracted using a Midi Prep kit (Qiagen). 3 kbp DNA was prepared from pGEM3z601 plasmid (3025 bp). The plasmid was grown and amplified in 100 μg/ml ampicillin. The plasmid was linearized using single-cut digestion by ScaI-HF restriction enzyme in 10× CutSmart Buffer (New England Biolabs, NEB). The samples were then purified using a PCR purification kit (Qiagen) followed by ethanol precipitation. 5 kbp linear DNA was prepared from pUC18-12x (5102 bp) plasmid. The pUC18-12x plasmid was grown and amplified in 100 μg/ml ampicillin. The plasmid was then extracted using a Qiagen Midi Prep kit and linearized using KpnI and EcoRI restriction enzymes in 10x CutSmart buffer (NEB), resulting in a final length of 5094 bp linear DNA. Two enzymes were chosen to linearize the plasmid to avoid self-ligation or cohesive ligation. The 10 kbp linear DNA was prepared from Cas9-pET28a (Addgene) plasmid by growing and amplifying in a 25 μg/ml kanamycin and extracted by Midi Prep kit (Qiagen). The linearization of the plasmid was done by digesting with KpnI-HF restriction enzyme in 10x CutSmart buffer (NEB), resulting in 9564 bp linear DNA. The purity of linearized plasmids was checked on a 1.5 % agarose gel run in 1x TAE buffer, at 90 V for 90 minutes (see Figure S1).

### Nanopore Experiments

The nanopores were made by pulling filamented quartz capillaries of OD = 1 mm and ID = 0.5 mm (QF-100-50-7.5, Sutter Instruments). First, the capillaries were cleaned using ethanol and then dried under nitrogen. Then the capillaries were pulled using Sutter Puller P-2000/G with the following two-line program: Line 1: heat (700), filament (3), velocity (35), delay (145), pull (75); line 2: heat (560), filament (0), velocity (15), delay (128), pull (200). The above program was optimized to get pore diameters in the range of 40-50 nm. The diameter of these pores was further reduced to the required (∼ 20 nm) size by e-beam exposure under a scanning electron microscope (SEM - Carl Zeiss Ultraplus FESEM) using a 3 keV electron beam^26,46^ (Figure S2). Nanopipettes (capillaries with the nanopore at their tip) were further cleaned in oxygen plasma in PlasmaPrep-III system (SPI supplies) for 1 minute at 50 W power of RF (at 250 mtorr pressure). This treatment also renders the nanopores hydrophilic. The capillaries were then attached to custom-made Teflon fluid cells with glue followed by filling of the fluid cell and the capillary with nanopore buffer (NPB, 4 M LiCl, 10 mM Tris-HCl, 1 mM EDTA, pH 8) (see schematic in Figure S3). The fluid cells were kept in a vacuum desiccator for 15 minutes to remove any air bubbles. After desiccation, the fluid cell was kept inside a Faraday cage to shield it from any external noises and connected to the current amplifier with Ag/AgCl electrodes. Voltage (300 - 900 mV, see text) was applied across the nanopore and the corresponding pore current was measured by Axopatch 200B amplifier (Molecular Devices). For each nanopore, linear I-V characteristics, low noise (< 5pA @ 5 kHz) and stable baseline current were ensured before adding any sample. Linear DNA was added to the fluid chamber (conc. 0.4 - 0.8 nM) and translocation events were recorded at the required voltage (see text). After recording ∼1000 translocation events, the polarity of the voltage bias was reversed to record reverse translocation data. Data acquisition was done at a sampling frequency of 200 kHz using NI DAQ PCI-6251 (National Instruments). The recorded data was filtered at 25 kHz for offline analysis. For recording and analysis of the data, we used custom-made LabVIEW programs (National Instruments). Note, all data comparison is done by measuring the samples on the same nanopore, back-to-back. Between sample changes, the fluid cell was rinsed with the nanopore buffer (NPB) multiple times (10x volume of fluid cell) and confirmed for no events and a stable baseline before adding the next sample.

## RESULTS AND DISCUSSION

### Characteristics of nanopore measurements

In Figure 1 we show the resistive pulse sensing technique of our glass nanopores. Figure 1a (left) shows SEM images of four typical nanopores with diameters ranging from 17 nm to 22 nm, fabricated at the tip of quartz capillaries. Optical image of the tapered capillaries is shown in Figure 1a (right). These nanopores are mounted on a custom-made fluid cell where the experimental buffer outside the capillary (cis chamber) and the buffer inside the capillary (trans chamber) are connected only through the nanopore (see schematic and picture of the nanopore setup in Figure 1a, 1b & S3). The two chambers were filled with nanopore buffer and connected to the amplifier. Upon applying a potential difference across the chambers, a constant ion flow was established, resulting in a baseline (open pore) current (Figure 1c). When DNA was introduced into the cis chamber, downward spikes (events) corresponding to ‘forward translocation’ of DNA from cis to trans chamber through the nanopore, were observed (from fluid cell, into the capillary, through the nanopore) (see schematic in Figure 1b (top) and Figure 1d). After collecting approximately 1000 forward translocation events, the polarity of the applied voltage was reversed. This reverses the open pore current and produced upward spikes during DNA translocation, indicative of ‘reverse translocation’ (from capillary into the fluid cell, through the nanopore) (see schematic in Figure 1b (bottom) and Figure 1e). From the current traces, translocation events for individual molecules were extracted and analysed for their conductance drops (ΔG) and translocation times (Δt) (see schematic in Figure 1f Top-right). Typical statistical distribution of the event parameters is shown in Figure 1f, as a scatter plot of the event conductance drops (ΔG) plotted against the event translocation times (Δt). The corresponding ΔG and Δt histograms are displayed in Figure 1f right and top respectively. The scatter plot shows three populations (marked with blue dotted circles). The population at low ΔG and Δt values is attributed to collision events and is typically filtered out^26,46,47^. Among the two main populations, the bottom one corresponds to unfolded DNA translocating in a linear conformation, while the upper one corresponds to events consisting of DNA translocating in fully or partially folded conformation^48^ (see Figure 2a). These are collectively referred to as ‘folded DNA’ events in this work. The ΔG histogram (Figure 1f, right) shows the unfolded and folded populations as two distinct peaks with mean values of 0.72 ± 0.04 nS (referred to as ΔG_1_) and 1.19 ± 0.7 nS (referred to as ΔG_2_), respectively. The Δt histograms are typically broad (due to a mix of folded and unfolded translocations) and are shown in Figure 1f (top). The mean translocation time of all the events was estimated as 0.87 ± 0.19 ms from the Gaussian fit. Note that the translocation events corresponding to multi-folded DNA (DNA folded multiple times) are also possible (for example, see Figure 2a), however, their occurrence is significantly rare for the DNA lengths considered in this study.

**Figure 1.**
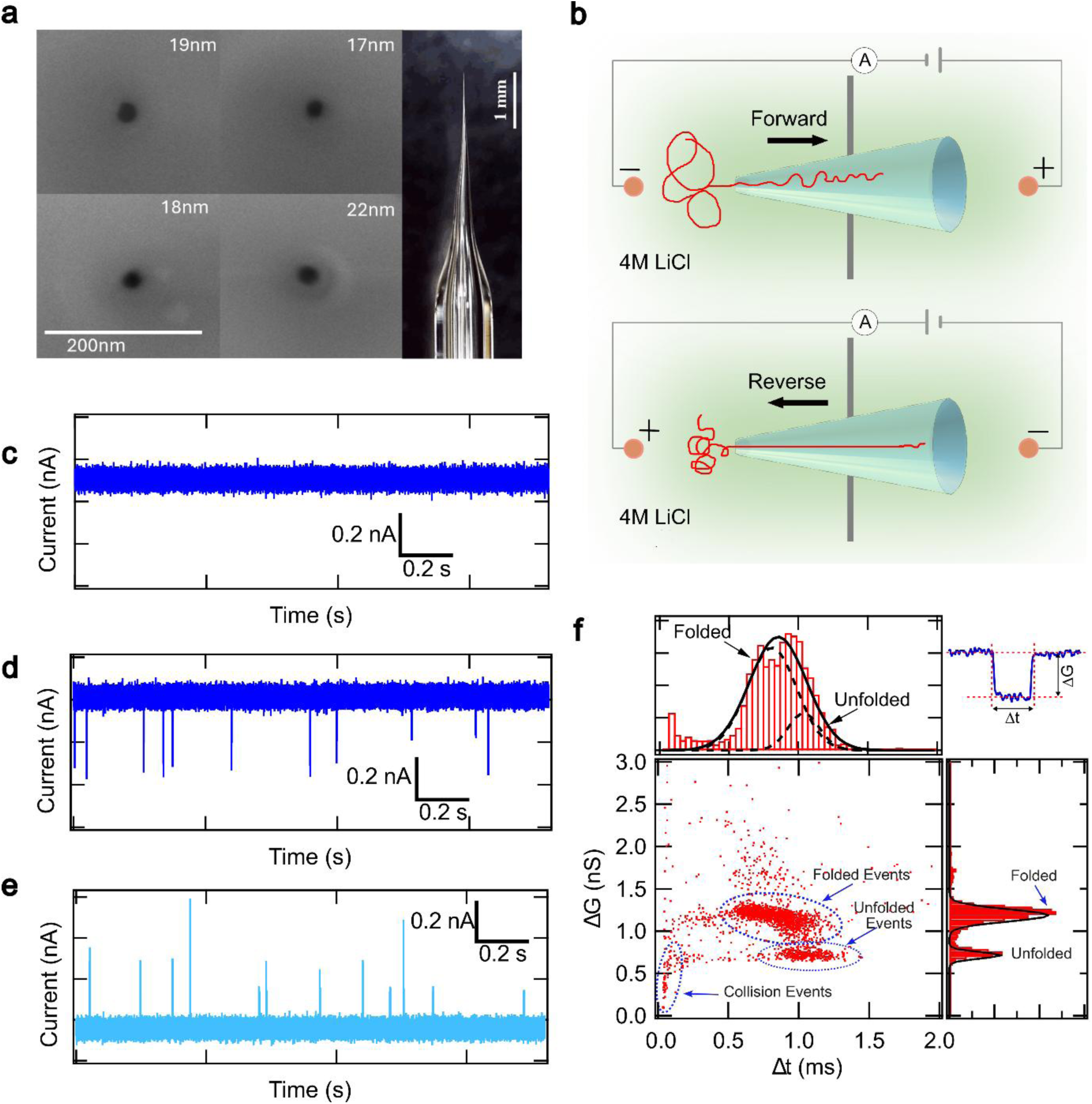
Characterization of nanopore device. (a) SEM images (left) of typical nanopores of ∼20 nm diameter and the side view of a capillary (right). (b) Schematic diagram of forward and reverse translocation of linear DNA through a nanopore with opposite voltage polarity. (c) The current through a nanopore with no sample added (16 nm pore in 4 M LiCl) and (d) & (e) show translocation events after adding DNA (10 kbp linear DNA, 300 mV voltage) in forward and reverse bias (−300 mV). (f) Scatter plot of the conductance drop (ΔG) against translocation time (Δt) and their respective histograms on the right and top, respectively. The multiple populations are due to collision events and different configurations of the DNA as marked by blue arrows. Gaussian fits to the isolated (dotted lines) and combined (solid lines) peaks in histograms are shown as black lines. Top-right shows a typical translocation event with the relevant parameters marked.

**Figure 2.**
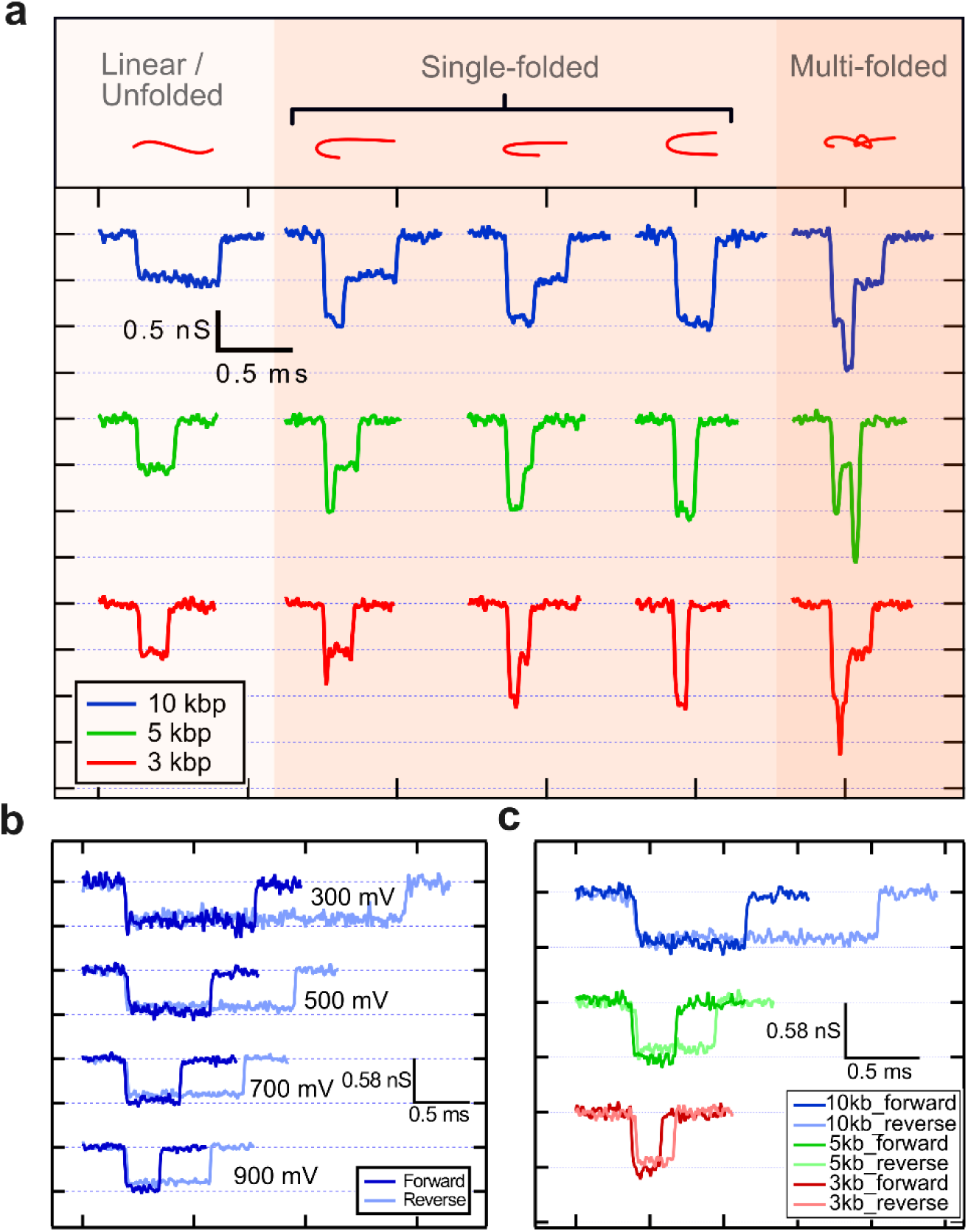
Library of events comparing different translocation possibilities. (a) The typical event signatures of translocation of DNA with different conformations (left to right) for three different DNA sizes (top to bottom), measured on the same nanopore at +500 mV. Possible DNA conformations for the event type are shown above in red. (b) Compares, on the same nanopore, forward and reverse DNA (10 kbp) translocation events of unfolded conformation at 8 different voltages, ± 300 mV, ± 500 mV, ± 700 mV, and ± 900 mV. The darker shades are for forward translocation (positive voltages), and the lighter shades are for reverse translocation (negative voltages). (c) shows representative events comparing forward and reverse translocation events (unfolded conformation)) on the same nanopore for varying DNA lengths (3, 5 and 10 kbp) at +500 mV. For all events, grid lines are chosen equal to mean value of ΔG_1_ histogram of respective datasets.

### Comparison of translocation events

In Figure 2, we present a sample of DNA translocation events in opposite directions for different DNA lengths and applied voltages. Figure 2a shows representative events of forward DNA translocation with increasing number of folds (left to right). Representative events are shown for 10 kbp (blue), 5 kbp (green), and 3 kbp (red) linear DNA, arranged from top to bottom, respectively. The multiple conductance levels observed in the events correspond to the multiple folds of DNA strands present in the pore during translocation^49,50^. Schematics illustrating the possible DNA conformations for each event type are shown above the respective events in red. The first column represents linear, unfolded events, while columns 2 to 4 show single-file translocation events with increasing degrees of folding. Column 5 represents higher-order folding, although such events are rare, as is evident from the frequency distribution in Figure 1f. The width of the event (translocation time (Δt)) corresponds to the length of the translocation DNA molecule.

To avoid confusion with events corresponding to various folded states, in the following, we isolate and compare only the unfolded events. The unfolded events are filtered by using a simple upper cut off, ΔG < ΔG_1_ + 2σ_1_. Here, ΔG_1_ and σ_1_ are the mean and the standard deviation, respectively, of the first peak in the ΔG histogram. In Figure 2b, we show the comparison of unfolded events during forward (dark lines) and reverse (lighter lines) translocation of 10 kbp DNA at applied voltages of ± 300 mV to ± 900 mV. From the data, it is evident that the translocation times are significantly longer in the reverse direction compared to the forward direction for all the voltages explored in this study. In Figure 2c, we show events comparing reversible translocation of different lengths of DNA measured at fixed voltage (± 500 mV). We note that the translocation time increases more significantly for the longer DNA lengths. This agrees well with the reports in the literature^34^. For every DNA length measured, a minimum of 1000 events were recorded at each voltage. This process was repeated eight times for each voltage: four in the forward direction (+300 mV, +500 mV, +700 mV, +900 mV) and four in the reverse direction (−300 mV, −500 mV, −700 mV, −900 mV). The nanopore was thoroughly washed with buffer multiple times between samples and an event-free stable baseline was confirmed to ensure the absence of any detectable residual DNA from previous samples. Since subtle geometrical variation in pore-to-pore fabrication may lead to differences in translocation dynamics, all sample comparisons drawn in this study were measured across the same pore, and the trends were reproduced in multiple nanopores (see Table S1).

In the following sections, we present a systematic comparison of key translocation parameters including ΔG, Δt, DNA folding fraction, and event charge deficit (ECD) as a function of translocation direction, DNA length, and applied voltage.

### Effect of voltage, DNA length, and direction of translocation on ΔG

Figure 3 shows ΔG histograms comparing conductance drops for different DNA lengths, translocation direction, and voltages. Figure 3a shows two peaks in the ΔG histograms. The first peak, ΔG_1_, corresponds to events when DNA is translocated in linear conformation and the second peak, ΔG_2_, corresponds to the translocation event of folded (partial or fully) DNA. This is seen for DNA of all lengths measured in this study (see Figure 3a inset). We also find that the ΔG_1_ and ΔG_2_ remain constant across DNA lengths. This is expected, as the conductance drop for a polymer longer than the pore length depends on its cross-sectional area and not on its length. Figure 3b compares ΔG_1_, ΔG_2_ in forward and reverse directions, measuring lower ΔG values in the reverse direction (∼10% for 5 kbp DNA at 500 mV). This reduction in ΔG values for reverse translocation is seen for other DNA lengths as well (see Figure 3b inset) and is also evident in the representative events shown in Figure 2b-c (see also Table S2). Finally, in Figure 3c we show the voltage dependence of ΔG values. We find that the ΔG values remain constant (within error bars) across the voltage range addressed in this study. We also find that the difference in ΔG values between translocation of forward and reverse direction, also remain consistent in this voltage range. This is shown in Figure 3c inset (solid lines represent forward translocation, and dotted lines represent reverse translocation). These measurements were repeated in multiple nanopores and additional datasets are shown in Figure S4.

**Figure 3.**
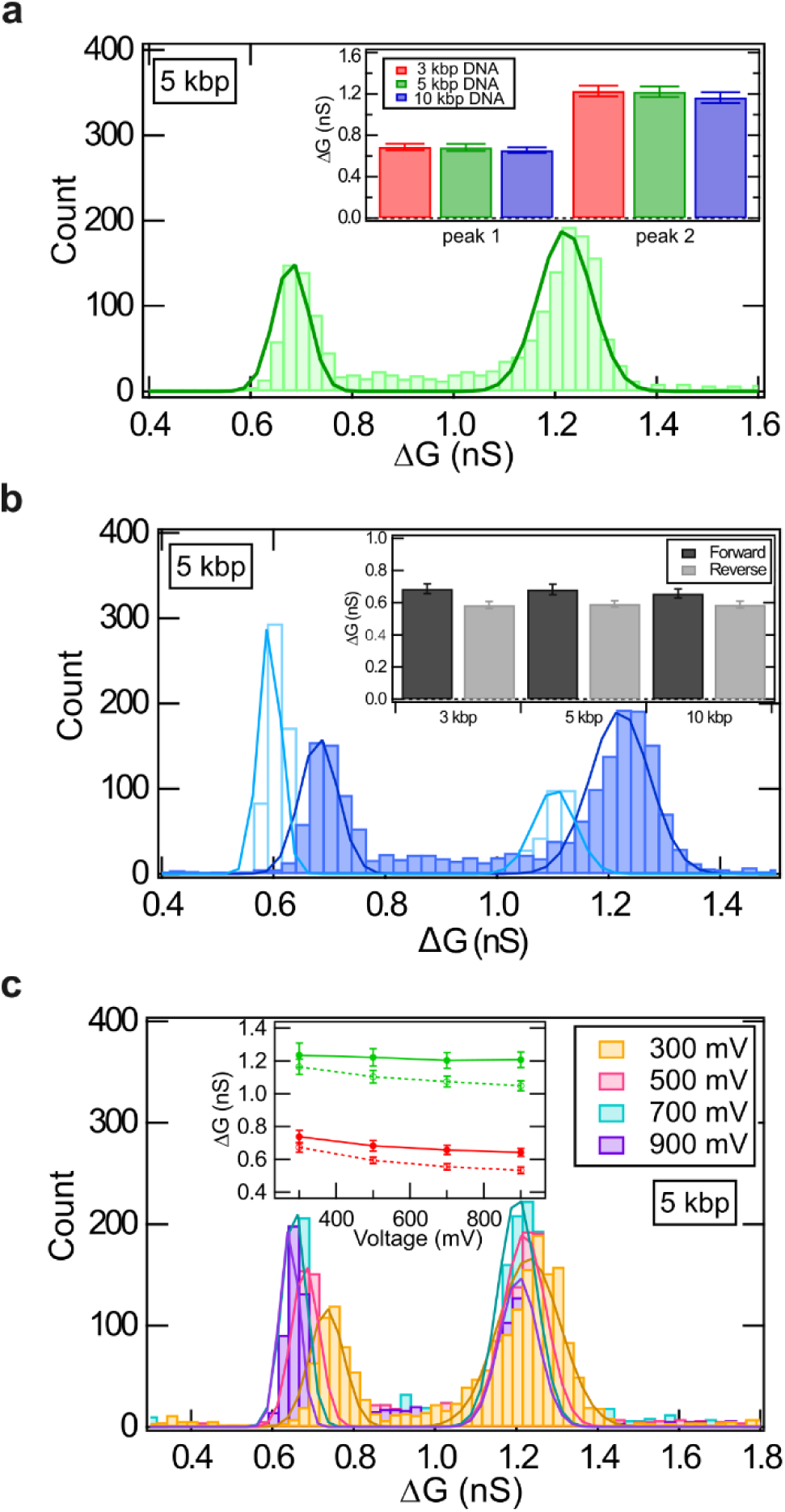
Effect of translocation direction on the conductance drop (ΔG) for different DNA lengths and applied voltages. (a) shows the two peaks in the ΔG histogram, corresponding to folded and linear conformations, for 5 kbp DNA at 500 mV. ΔG_1_ and ΔG_2_ values for 10 kbp, 5 kbp, and 3 kbp DNA are compared in the inset. (b) compares ΔG histograms for forward (+500 mV, dark) and reverse (−500 mV, lighter shade) translocation of 5 kbp DNA. The same comparison for other DNA lengths is shown in the inset. (c) ΔG histograms of 5 kbp linear DNA translocation measured at +300, +500, +700 & +900 mV voltages. Inset shows the mean values of ΔG with both forward and reverse translocation of 5 kbp DNA with varying voltages. The error bars represent the standard deviations of the Gaussian fits.

Note, since the changes in ΔG histograms shown in Figure 3 are small, we reconfirm these changes by calculating ΔG values from the analysis of G-histograms. All the collected raw event traces, measured as G (conductance) versus time, are concatenated and their histograms are plotted. These histograms have peaks corresponding to the baseline (G_0_), linear DNA level (G_1_) and folded DNA level (G_2_) (see Figure S5). All-point histograms of these traces are analyzed as G histograms. Peaks in G histograms were fitted with Gaussian distributions and ΔG_1_ and ΔG_2_ are calculated as ΔG_1_ = G_1_ – G_0_ & ΔG_2_ = G_2_ – G_0_, respectively. These values corroborate our results in Figure 3. Our results on reduced ΔG values for reverse translocation agree well with the literature on conical nanopores^32^. This difference in ΔG values of the forward and reverse events is attributed to bias-dependent redistribution of charges at the conical wall of glass nanopores.

### Translocation time analysis

We next show a detailed comparison of the statistics of the event translocation times (Δt). Figure 4a shows the Δt histograms for three different DNA lengths at +500 mV, measured back-to-back on the same nanopore. The peak values from Gaussian fits were found to be 0.56 ± 0.12 ms, 0.31 ± 0.06 ms, and 0.23 ± 0.05 ms, for 10 kbp, 5 kbp and 3 kbp DNA, respectively. We see longer translocation times for longer DNA lengths, as expected^34,51,52^. Since the translocation times of the DNA can vary depending on its various folded conformations, for further analysis, we isolate and compare translocation times (Δt_unfolded_) of events that result only from the linear conformation.

**Figure 4.**
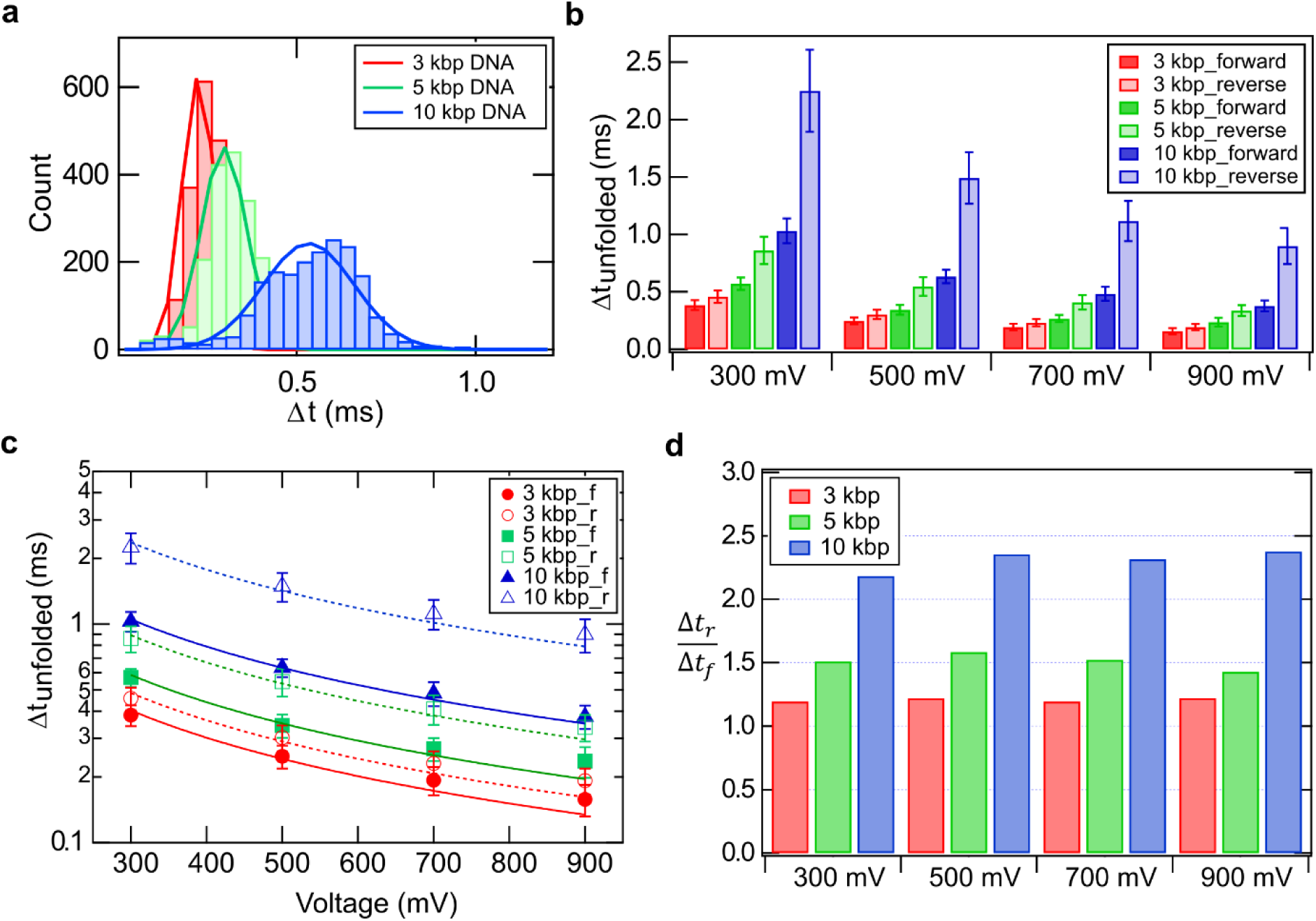
Analysis of Translocation times (Δt) with varying DNA lengths, applied voltage and translocation direction. (a) Comparison of Δt histograms for both linear and folded events of 3 kbp (red),5 kbp (green) and 10 kbp DNA (blue), at 500 mV. (b) Bar plots comparing the mean translocation times of unfolded DNA events with respect to voltage and translocation direction. The error bars represent the standard deviation of the Gaussian peak. In (c) we plot the translocation times of the unfolded population as a function of applied voltage. The data is fitted to Δt ∼ 1/V_applied_. Solid and dotted lines are fits for forward and reverse translocations, respectively. The error bars are the standard deviation of the Gaussian fits. Fit values are given in Table S3. (d) The ratio of the translocation times (unfolded population) in the reverse to the forward direction for different applied voltages and DNA lengths. The values of the ratio are given in Table S4.

In Figure 4b, we compare the mean values of the Δt_unfolded_ histograms (as bar plots) for different DNA lengths measured at different voltages. Data is shown for 3 kbp DNA (red), 5 kbp DNA (green), and 10 kbp DNA (blue). For the forward direction, solid bars are used, while bars with lighter shades are used for the measurements in the reverse direction. We note that, for each voltage, the Δt_unfolded_ is maximum at 300 mV, for all DNA lengths, and then decreases monotonically with increasing voltage. This voltage dependence is seen for both forward and reverse translocations. We show this voltage dependence explicitly in Figure 4c for each DNA length. This change in translocation times was found to depend inversely on the applied voltage. This is seen by fits to Δt = C/V_applied_ (where C is the fitting parameter), shown in Figure 4c. The error bars represent the standard deviations of the Gaussian fit. Since, for ∼ 20 nm nanopores, the DNA translocation is primarily electrophoretic in nature, the reduction in translocation times with increasing voltage is expected and its inverse voltage dependence agrees well with the literature^53–56^ (See Figure S6 & S7 for additional datasets). We observed that the proportionality constants (per basepair) are similar for the three DNA lengths measured (see Table S3).

We also see that the unfolded DNA translocation time, Δt_unfolded_, is longer for reverse translocation when compared to the forward for all given voltage and DNA length conditions. The ratio of Δt_unfolded_, ((Δt_r_)/(Δt_f_)), for the reverse translocations to the forward translocations is plotted in Figure 4d. For a given DNA length, this ratio remains constant across all voltages; however, at a fixed voltage, the ratio increases with DNA length (Table S4). This trend suggests that the relative increase in Δt during reverse translocation is inherently dependent on interactions that depend on the DNA polymer length and is independent of the applied electric field.

The slowing down of DNA in reverse translocation could be due to multiple reasons. One of the arguments proposed is that during forward translocation, the DNA buckles and faces less drag force while passing through the pore, while in reverse motion the DNA is under tension, which results in polymer stretching leading to a longer effective length of the DNA. This may result in a higher drag force and hence DNA slows down^34^. An alternate hypothesis could be that, as the DNA polymer approaches the nanopore from the reverse direction, its conical geometry causes the DNA to elongate and interact more extensively with the sloped pore walls. These interactions generate additional frictional resistance during translocation. This frictional resistance would be higher for longer DNA, which is seen in Figure 4d. The constant slope of the ((Δt_r_)/(Δt_f_)) indicates that the increase in reverse translocation times with DNA length remains constant across all applied voltages. This suggests primarily size-dependent interaction with the nanopore walls rather than an electric field-dependent one. This highlights the significance of both pore wall interactions and DNA length in modulating translocation dynamics through a nanopore.

### Quantifying folded conformations during DNA translocation

We next analyze the folding characteristics of DNA, during translocation through the nanopore, by examining their electrical signatures. Figure 5a shows the scatter plot of 3 kbp and 10 kbp DNA at ± 300 mV in both forward and reverse directions. It shows that the population of events corresponding to linear and folded configurations, of each DNA length, are vertically separated (along the ΔG axis) but overlap horizontally (along the Δt axis). For shorter DNA lengths the forward and reverse populations are indistinguishable but the contrast increases for longer DNA lengths.

**Figure 5.**
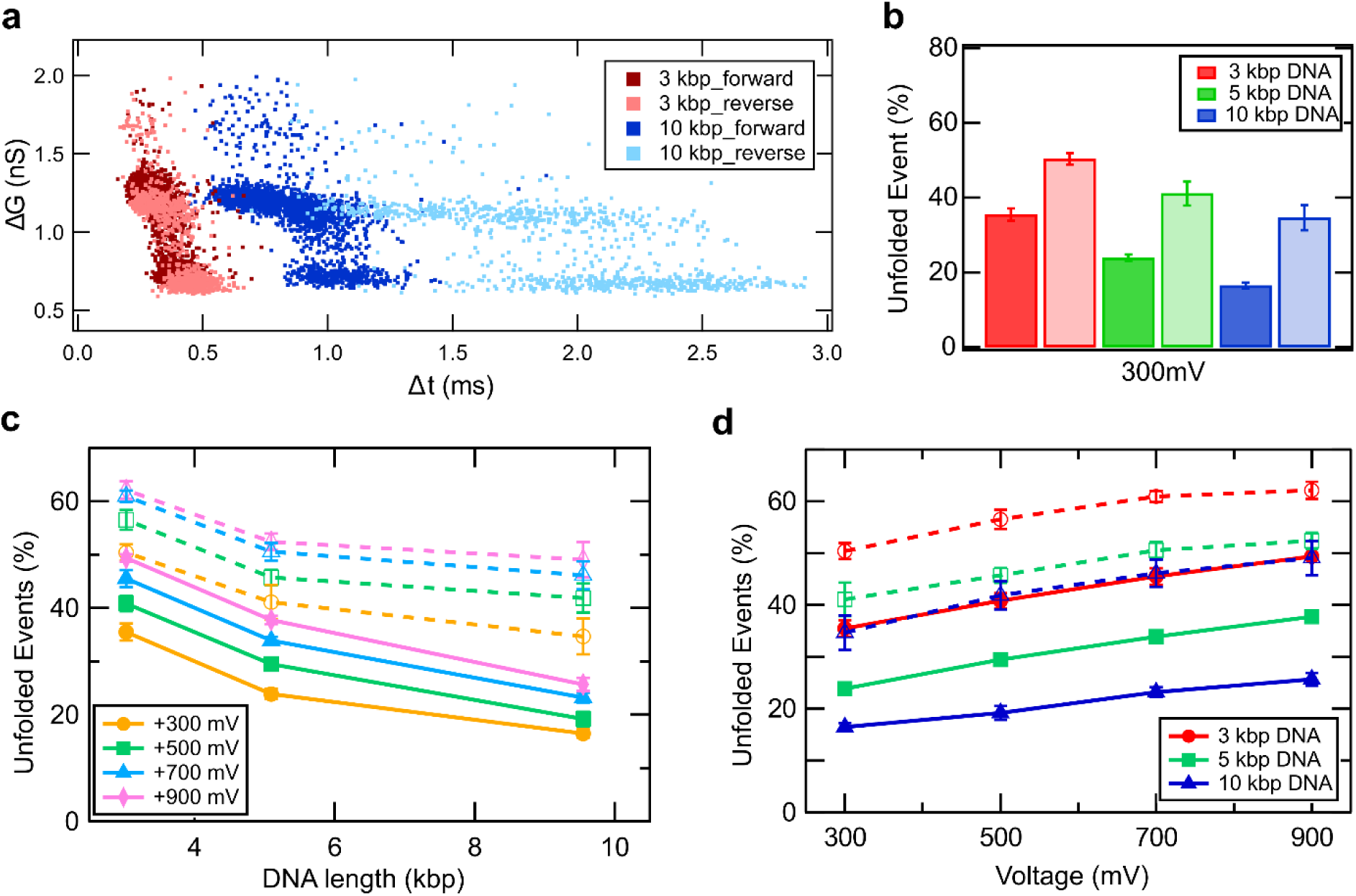
Statistical analysis of DNA translocation in folded conformation. (a) shows scatter plot of 3 kbp and 10 kbp DNA in forward (+300 mV, dark shade with 1917 and 2109 events, respectively) and reverse direction (−300 mV, lighter shade with 1199 & 947 events respectively). In each direction, folded and unfolded translocation events are measured as distinct population clusters. Scatter plot for 5 kbp DNA is not included, for visual clarity. (b) compares percentage of unfolded events, as bar plot, during forward (dark shaded bars) and reverse (lighter shaded bars) translocations for the 3 different DNA lengths at +300 mV. The length and voltage dependence of unfolded events percentage, is shown in (c) and (d), respectively. Here the solid and dotted lines joining the data points correspond to forward and reverse translocations, respectively. The data trends presented here are reproduced and averaged over all the nanopores used in this study (see Table S5 and S6).

We note that, in our experimental conditions, almost the entirety of the events recorded are either linear or single-folded. Higher folded conformations do appear but are very rare for the DNA lengths considered^57^ (see Figure 2a). By counting the number of events in each cluster, we calculate the percentage of molecules that translocated in unfolded (linear) configuration. We quantify these percentages of unfolded events as a function of translocation direction (Figure 5b), DNA length (Figure 5c) and the applied voltage (Figure 5d). In Figure 5b we show that during forward translocation of 3 kbp DNA, up to 33 % of the DNA molecules translocate in unfolded conformation. This proportion decreases monotonically with increasing DNA length, and drops to less than 20 % for 10 kbp DNA. In contrast, for reverse translocation at – 300 mV, more than 50 % of the molecules translocate in the linear fashion. Similar to forward translocations, the percentage of unfolded population in reverse translocations decreases with DNA length (see Figure 5c). This indicates a higher tendency of longer DNA to translocate in folded conformations. Additionally, the proportion of unfolded translocations increases with applied voltage across all DNA lengths studied (Figure 5d). This is consistent for all DNA lengths explored in this study and the results are reproduced with multiple nanopores (see Figure S8, S9 & Table S5 & S6).

The results for forward translocation can be understood from the fact that longer DNA has statistically larger propensity of folding and hence shows lower unfolded event percentage. Further, in reverse translocation, we see more unfolded events. This is in contrast with some recent studies done with longer DNA^33^. In our understanding, the enhanced occurrence of unfolded events, seen in our experiments with 3 kbp – 10 kbp DNA range, may be attributed to the conical geometry of the nanopore. In reverse translocation, the DNA molecules are electrophoretically funnelled towards the pore through the tapered capillary. The surface interactions due to the confining geometry along the pore wall promote unfolding of the DNA polymer^58^. This may lead to our reported higher fraction of unfolded translocation events. In contrast, during the forward direction, DNA is captured in the nanopore directly from the bulk solution, and no such gradual wall confinement is available. Furthermore, working at higher voltages may result in more unfolding of DNA, which is corroborated by the slight increase in unfolded event percentage seen at higher voltages (Figure 5d).

### ECD analysis of DNA translocation dynamics

The translocation parameters discussed in previous sections depend explicitly on the conformation of the translocating polymer (DNA). We analyse the event charge deficit (ECD) of our translocation data which characterizes the volume of the translocating DNA and is independent of the folded configurations of the polymer. We first show a library of event types for 10 kbp DNA with different levels of folded conformations in both forward (Figure 6a) and reverse (Figure 6b) directions. The possible DNA conformations, corresponding to distinct levels in the events, are shown above individual events in red. The ECD of an event is defined as the net charges excluded by the analyte molecule during its translocation through the nanopore and is evaluated as 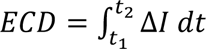, where t_1_ and t_2_ are the start and end of the event^46,57,59^. This, numerical integration of the area under the event, is schematically shown in the inset of Figure 6c. For the events of both forward and reverse directions, Figure 6c and 6d show the corresponding ECD histograms. Notably, despite the variation in DNA folding states, all events for a given DNA length yield a single peak in the ECD histograms. This indicates that the ECD value is independent of the folded conformation of DNA during translocation and depends only on the total DNA volume. We plot ECD histograms for the different DNA lengths in both directions in Figure 6e. Firstly, we note that the mean ECD values scale with DNA length. In forward direction, the longer 10 kbp DNA has ECD value of 1003 ± 80 ke as compared to 378 ± 43 ke for 3 kbp DNA at +300 mV. Given that the DNA diameter is same for all, ECD scales almost linearly (see Figure 6e inset) with the range of DNA lengths studied in this work.

**Figure 6.**
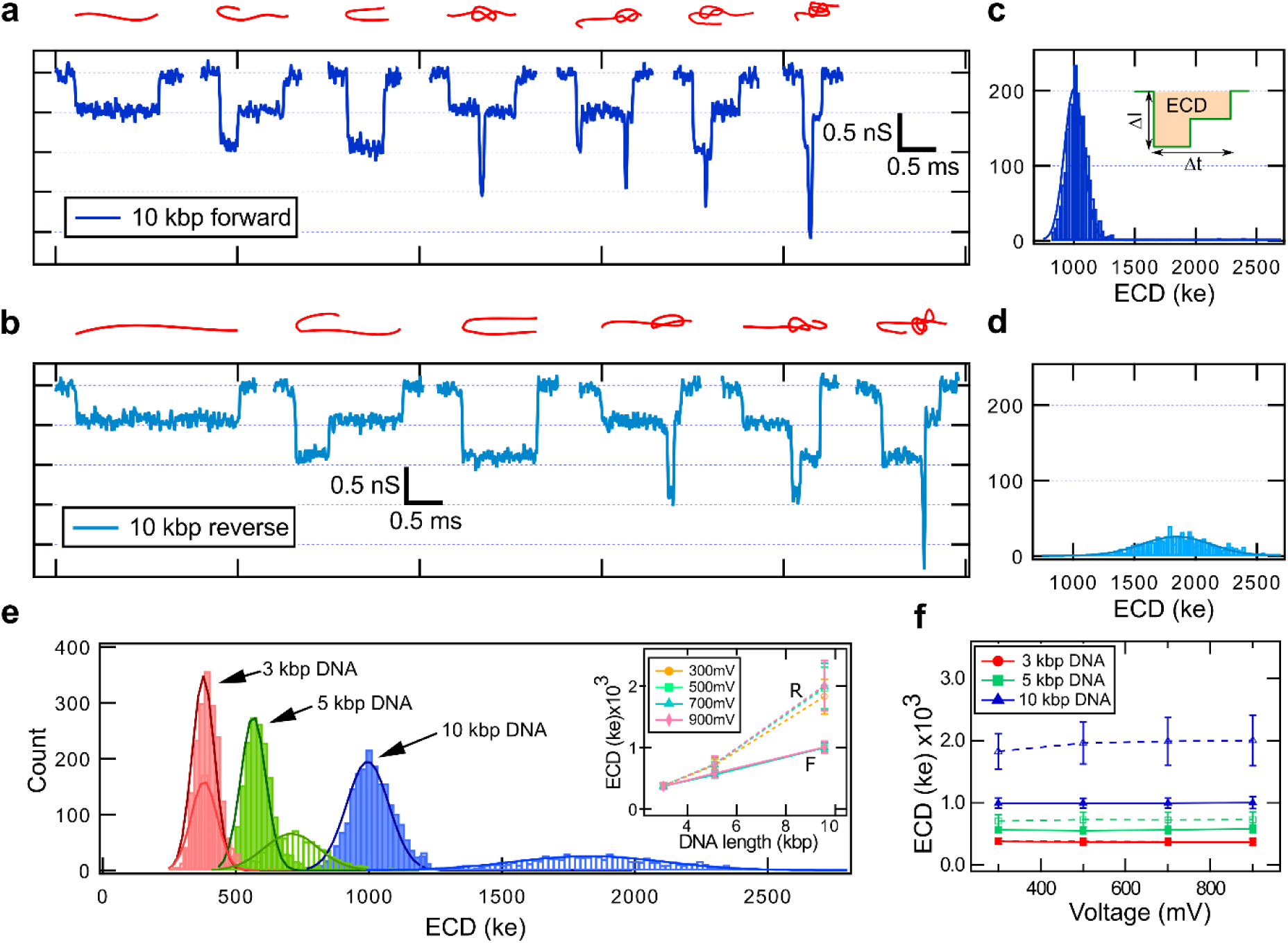
ECD analysis of translocation dynamics. Representative events for (a) forward (+300 mV) and (b) reverse (−300 mV) translocation of 10 kbp DNA showing different folds seen as distinct conductance levels in the event. The possible DNA conformations corresponding to the levels are shown in red on top of each event. (c) & (d) shows ECD histograms of the datasets shown in (a) and (b) respectively. (c) inset shows the schematic for ECD estimation (shaded region) for a typical event (green). (e) compares ECD histograms (300 mV) of 3 kbp, 5 kbp & 10 kbp DNA for translocation in forward (filled bars) and reverse directions (open bars). Inset shows dependence of ECD values on DNA lengths and applied voltages (f). Solid lines are for forward (marked F) and dashed lines are for reverse (marked R) translocations. The error bars represent the standard deviation of the Gaussian fit to ECD histograms.

During reverse translocations, the ECD histograms are also found to increase with DNA length. For shorter DNA (3 kbp), the mean ECD values are comparable between forward and reverse translocations. However, for longer DNA (5 kbp and 10 kbp), we observe that reverse translocation yields a higher mean ECD and larger standard deviation compared to the forward case. This difference increases with DNA length (see Figure 6e (inset)). Next, we tabulate the ECD ratios for reverse and forward translocation in Table S7. Finally, in Figure 6f we show the voltage dependence of ECD. We find that for all DNA lengths and for both translocation directions, the mean ECD remains largely unaffected by the applied voltage. This trend is consistent with the counteracting effects of increasing current blockage (ΔI) and decreasing dwell time (Δt) with increasing voltages, resulting in an overall voltage-independent ECD.

It is important to note that both ΔG and the ECD parameters exhibit dependence on the DNA size. However, they capture different aspects of the translocation process. The ΔG parameter is sensitive to the DNA folding conformations, making it more suitable for studying folding statistics. In contrast, ECD is independent of the polymer folding and is a suitable parameter for precise size estimation of an unknown DNA. These findings highlight the distinct but complementary roles of ΔG and ECD in nanopore-based DNA characterization. A combination of the two parameters can be used to further quantify the conformation states of the translocating polymer as well as structural details of the nanopore itself (see section below).

### ECD analysis to quantify DNA fold lengths and nanopore sensing length

Finally, we demonstrate that the ECD analysis is powerful enough that it can be leveraged not only to quantify fold lengths of the translocating DNA molecule but also to in-situ calibrate the sensing region of the conical nanopores. We first show DNA fold lengths quantification from nanopore measurements. Figure 7a displays several events for 10 kbp DNA, with varying folded lengths, translocating through the nanopore. The folded regions were identified from the current data by threshold analysis as detailed in our previous work^46^ and are shown in red. Note, for this analysis, only those events are considered where the DNA is folded only once. The length of the folded region, the ‘fold length (L_f_)’, was calculated using the established linear relationship between ECD and DNA length, expressed as^4,46,57^,

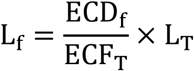

where ECD_T_ is the total ECD of the event, L_T_ is the total length of the DNA (for example, for our 10 kbp sample, L_T_ = 9546 bp = 3245.64 nm), and ECD_f_ is the ECD of the folded part (red region) of the event. This allows for precise measurement of the folded part of the DNA. We quantify L_f_ for all folded events and the resulting fold length histogram is shown in Figure S10a. We note that shorter fold lengths occur more frequently and the maximum fold length observed is half of the DNA’s total length, as expected. Additionally, we note that the current level corresponding to the folded configuration appears at the beginning of most events. This is confirmed by checking the start-position of the folded region with respect to the start of the event (shown in Figure S10b). This implies that predominantly folded end of the DNA enters the pore first^50,60^.

**Figure 7:**
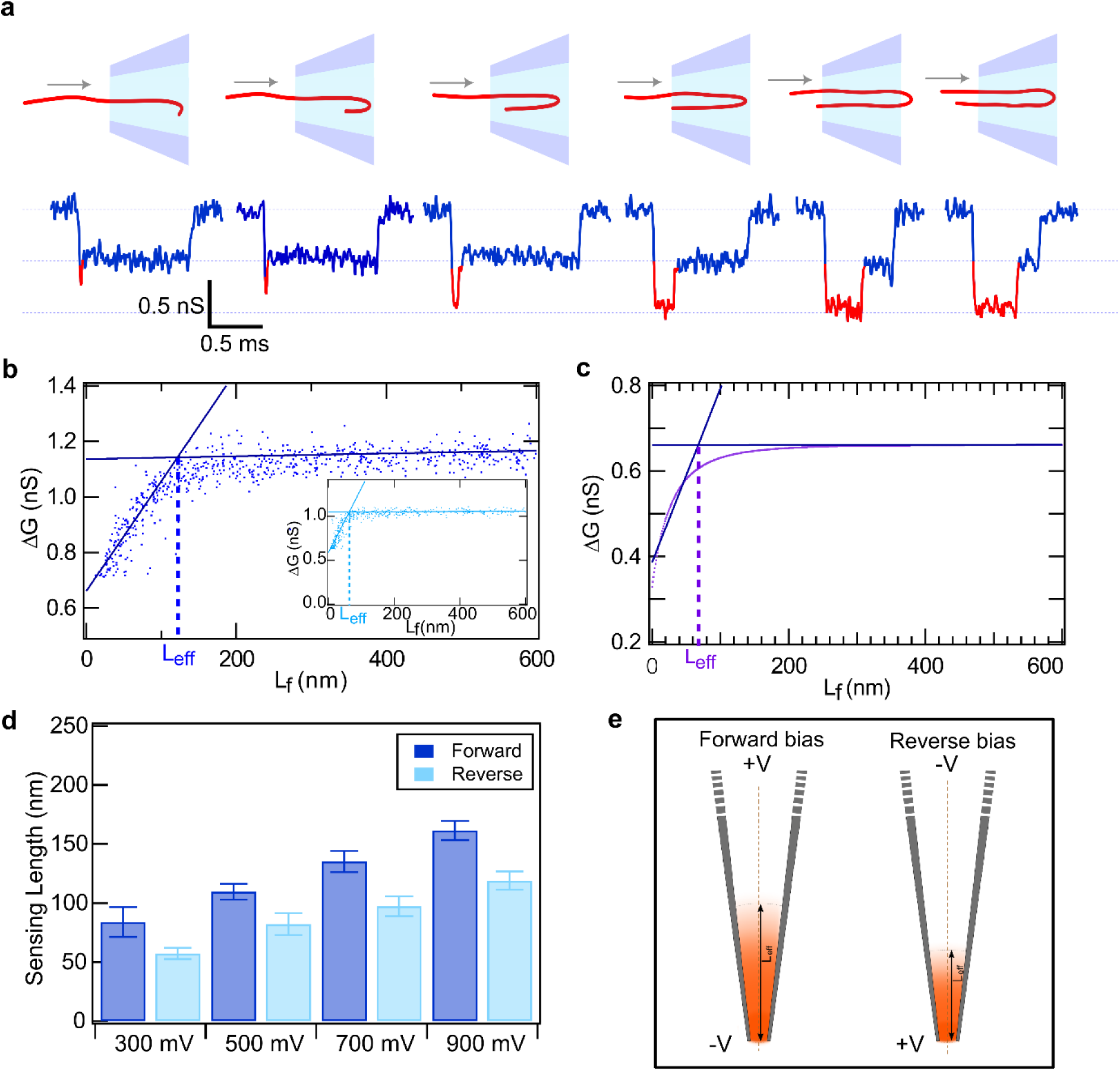
Determination of sensing length of a nanopore by ECD analysis. (a) highlights a few events of 10 kbp DNA (16 nm nanopore) where the folded region of the DNA is highlighted in red. From left to right, we show events with increasing lengths of folded region (L_f_). (b) Scatter plot of ΔG vs L_f_ for 10 kbp DNA at +500 mV (−500 mV in inset). The data is fitted with two piecewise linear functions. Estimated L_eff_ values for forward and reverse directions are shown using dotted lines. (c) Simulated plot of ΔG vs L_f_ and corresponding L_eff_ is shown by dotted line. (d) Bar plots to compare the normalized L_eff_ values between forward and reverse directions at indicated voltages. The data is averaged from multiple (N = 7) experiments (See Figures S11-S13 for individual data). (f) Schematic diagram showing sensing length as the high field region in both forward (left) and reverse (right) directions.

Further, we note that for smaller L_f_ values, the event depth (ΔG) increases with L_f_ (as seen in the first three events of Figure 7a) and then saturates to the folded DNA level (last three events in Figure 7a). We plot L_f_ values with their corresponding ΔG values for all the folded events as a scatter plot in Figure 7b. We clearly see that ΔG linearly increases with L_f_ at smaller values and then transitions to a constant value at large L_f_. This behaviour is consistent for all the DNA lengths used in this study (see Figure S10c). The transition between the two slopes is clearest in the 10 kbp data and the transition point is identified by fitting a piecewise linear function (solid lines in Figure 7b, see Figure S10d) to the two regions, and is labelled as L_eff_ (shown as vertical dotted line). We propose that L_eff_ represents the sensing length of the nanopore. DNA molecule, when inside the nanopore, causes a conductance drop to a level that corresponds to the DNA cross-section. DNA with a small folded region adds an additional conductance drop (ΔG), measured as a spike in the event on top of the DNA level (see first event in Figure 7a). When DNA with a longer folded region is in the pore, it causes a greater drop in the conductance (see second event in Figure 7a). This continues until the folded region becomes equal to the effective length of the nanopore or its sensing length (see schematic cartoon in Figure 7a). At this point spike height has reached 1-fold DNA level. For DNA with fold lengths (L_f_) longer than the nanopore sensing length (L_eff_), folded DNA spikes remain the same height but increase in width (see last three events in Figure 7a). The increasing folded lengths of the translocating DNA transitions in the nanopore signal from increasing spike heights to increasing spike widths (See Figure S11 & S12 for additional datasets).

Thus, the DNA fold length (L_f_), at which the ΔG transitions from monotonically increasing to saturated phase, gives us a quantitative estimate of the effective sensing length of the nanopore. To confirm this geometric reasoning of the data in Figure 7b, we employed a double-cone model^43,45^ of our nanopore to calculate the pore conductance with and without the DNA molecule inside it (see Figure S14). In the conical geometry model, the ΔG corresponds to the change in conductance caused by the presence of DNA volume inside the pore (cone volume). The larger the DNA length in the pore, the higher the ΔG values. The sensing length of the conical nanopore, L_eff_, is the cone length when the pore volume is large enough that further increasing linear DNA volume causes no further change in pore current, saturating the ΔG. We used it to calculate conductance drops for different DNA fold lengths, and the result is shown in Figure 7c. We find that the simple geometric model captures all the features of the experiment, yielding a sensing length of 71 nm which is in excellent agreement with our experimentally measured range of L_eff_ (50 – 150 nm, Table S8 & S9) for different nanopores of similar diameter.

Although the geometric model shows all the essential features of our experiment, it is also independent of the applied voltage or the direction of DNA translocation. In our further experiments, we see that for forward translocations the L_eff_ monotonically increases with the applied voltage (Figure 7d). This implies that the sensing length of the nanopore is further constrained by the range of electric field distribution inside the conical nanopore. The range of the electric field distribution should increase with the applied voltage. We see the same trend of increasing L_eff_ with voltage in Figure 7d (See Figure S13 for data reproducibility). This was found true for both forward and reverse translocation directions. Inside a conical pore, it is known that the range of electric field distribution in reverse bias is smaller than the forward bias, due to ion concentration polarization caused by pore asymmetry^40^. This is seen in our data as the L_eff_ measured for reverse translocations (light bars, Figure 7d) is smaller than the L_eff_ measured during forward translocations (dark bars in Figure 7d). These features of our model are shown in Figure 7e.

## CONCLUSIONS

In summary, we present a comprehensive experimental investigation into the influence of confining geometry on electrophoretic movement of DNA by studying bidirectional translocation of linear DNA through a conical glass nanopore. By systematically tuning the bias voltage and analyzing the conductance drops, translocation times, DNA folding statistics, event charge deficit (ECD), and the nanopore sensing lengths, we evaluate how DNA of varying lengths (3–10 kbp) interrogates the conical pore geometry during translocation.

Our findings reveal that the conductance drop during reverse translocation reduces by ∼10 % when compared with the forward direction. Significant changes are seen in translocation times of the DNA through the pore. We show that the DNA translocates slower in the reverse direction and the effect gets more pronounced with increasing DNA length. When averaged over all the voltages addressed in this study, 3 kbp DNA reverse translocation times are 1.1 times slower than the forward translocations which increases to 1.8 times for the longer 10 kbp DNA. This suggests enhanced wall interactions for longer molecules. This wall-dependent slowdown is accompanied by a notable decrease in DNA folding during reverse translocation. For example, at 300 mV, the unfolded translocation fraction increased by 1.42x (for 3 kbp) to 2.1x (for 10 kbp DNA). Also, we show that independent of the translocation direction, the unfolded fraction increases with the applied voltage. This is particularly advantageous for potential applications in screening for DNA-protein interactions, where folded DNA could interfere with protein position mapping.

The sensing length of a conical nanopore has been a particularly elusive quantity to measure. We employ ECD-based analysis to calculate DNA fold lengths, providing an essential metric for direct estimation of the nanopore’s sensing length. We show that the sensing length of the conical nanopore depends on the asymmetry of the nanopore and the applied voltage. A numerical model was developed to verify our experiments, with results demonstrating close agreement with data. Our experiments reveal typical sensing lengths of 84 – 161 nm for forward biased (300 to 900 mV) nanopores, which reduces to 57 – 119 nm range in reverse biased conical nanopores. From the literature, we attribute this difference in sensing lengths to the electric field reconfiguration caused by ion concentration polarization during forward and reverse biases. We note that the sensing length increases with voltage, which may be due to voltage-driven extension of the high-field region inside the conical nanopore.

In conclusion, this work provides critical insights into how conical nanopore geometry and voltage polarity influence DNA translocation, an area of significant importance for the advancement of nanopore-based biosensing technologies. By systematically characterizing bidirectional DNA movement and developing a robust method to estimate the sensing length using event charge deficit (ECD), we offer a detailed framework for understanding the spatial resolution of asymmetric nanopores. Our findings reveal clear directional dependence in translocation dynamics in key translocation parameters, including conductance drops, translocation times, folding behaviour, and variations in sensing length. These effects help in understanding the role of pore geometry and electric field asymmetry in shaping molecule– nanopore interactions under a nanoscale confinement. The implications of these insights are particularly relevant for single-molecule applications that demand high spatial precision, such as mapping DNA–protein interactions, localization of protein or biomarkers on DNA, epigenetic profiling etc. Moreover, our ECD-based sensing length estimation offers a broadly applicable, *in-situ* & optics-free approach to assess and compare the sensing performance of diverse nanopore architectures. Altogether, this study delivers both conceptual and practical contributions toward engineering next-generation nanopore platforms with enhanced analytical sensitivity and spatial resolution.

## SUPPORTING INFORMATION

Contains tabulated data quantifying the histograms in the main text, gel images of purified linear DNA, picture of the experimental setup, SEM images of the nanopores used in this study, reproducibility data of nanopore experiments on multiple nanopores. Supplementary file also contains details of the model used to estimate sensing length. The file contains supplementary tables (S1-S9) and figures (S1-S14) as mentioned in the text.

## ACKNOWLEDGEMENTS

We acknowledge RRI’s internal funding and assistance from Mr. Yatheendran & RRI SEM facility and workshop in this work. We also acknowledge Ms. Divya Shet for assistance in sample preparation.

## AUTHORSHIP CONTRIBUTIONS

GVS, and SS conceived, planned the experimental protocols. SS contributed in sample preparation and performed experiments. SS and GVS performed data analysis. All authors contributed in manuscript writing.

## DISCLOSURE OF CONFLICTS OF INTEREST

No relevant conflicts of interest to declare.

## Notes

### Competing Interest Statement

The authors have declared no competing interest.

## REFERENCES

(1) Chen, K.; Choudhary, A.; Sandler, S. E.; Maffeo, C.; Ducati, C.; Aksimentiev, A.; Keyser, U. F. Super-Resolution Detection of DNA Nanostructures Using a Nanopore. Advanced Materials 2023, 35 (12), 2207434. 10.1002/adma.202207434.

(2) Ying, Y.-L.; Hu, Z.-L.; Zhang, S.; Qing, Y.; Fragasso, A.; Maglia, G.; Meller, A.; Bayley, H.; Dekker, C.; Long, Y.-T. Nanopore-Based Technologies beyond DNA Sequencing. Nat. Nanotechnol. 2022, 17 (11), 1136–1146. 10.1038/s41565-022-01193-2.

(3) Confined Nanopipette Sensing: From Single Molecules, Single Nanoparticles, to Single Cells - Yu - 2019 - Angewandte Chemie International Edition - Wiley Online Library. https://onlinelibrary.wiley.com/doi/full/10.1002/anie.201803229 (accessed 2025-07-08).

(4) Yang, W.; Restrepo-Pérez, L.; Bengtson, M.; Heerema, S. J.; Birnie, A.; der Torre, J. van; Dekker, C. Detection of CRISPR-dCas9 on DNA With Solid-State Nanopores. Nano Letters 2018. 10.1021/acs.nanolett.8b02968.

(5) Kowalczyk, S. W.; Hall, A. R.; Dekker, C. Detection of Local Protein Structures along DNA Using Solid-State Nanopores. Nano Lett. 2010, 10 (1), 324–328. 10.1021/nl903631m.

(6) Ratinho, L.; Meyer, N.; Greive, S.; Cressiot, B.; Pelta, J. Nanopore Sensing of Protein and Peptide Conformation for Point-of-Care Applications. Nat Commun 2025, 16 (1), 3211. 10.1038/s41467-025-58509-8.

(7) Chau, C. C. C.; Weckman, N. E.; Thomson, E. E.; Actis, P. Solid-State Nanopore Real-Time Assay for Monitoring Cas9 Endonuclease Reactivity. ACS Nano 2025, 19 (3), 3839–3851. 10.1021/acsnano.4c15173.

(8) Prajapati, J. D.; Pangeni, S.; Aksoyoglu, M. A.; Winterhalter, M.; Kleinekathöfer, U. Changes in Salt Concentration Modify the Translocation of Neutral Molecules Through a ΔCymA Nanopore in a Non-Monotonic Manner. Acs Nano 2022. 10.1021/acsnano.1c11471.

(9) Electrochemical Monitoring of Real-Time Vesicle Dynamics Induced by Tau in a Confined Nanopipette - Chen - 2024 - Angewandte Chemie International Edition - Wiley Online Library. https://onlinelibrary.wiley.com/doi/full/10.1002/anie.202406677 (accessed 2025-07-08).

(10) Oshidari, R.; Strecker, J.; C. Chung, D. K.; Abraham, K. J.; Chan, J. N. Y.; Damaren, C. J.; Mekhail, K. Nuclear Microtubule Filaments Mediate Non-Linear Directional Motion of Chromatin and Promote DNA Repair. Nature Communications 2018. 10.1038/s41467-018-05009-7.

(11) Ying, Y.-L.; Hu, Z.-L.; Zhang, S.; Qing, Y.; Fragasso, A.; Maglia, G.; Meller, A.; Bayley, H.; Dekker, C.; Long, Y.-T. Nanopore-Based Technologies beyond DNA Sequencing. Nat. Nanotechnol. 2022, 17 (11), 1136–1146. 10.1038/s41565-022-01193-2.

(12) Ratinho, L.; Bacri, L.; Thiebot, B.; Cressiot, B.; Pelta, J. Identification and Detection of a Peptide Biomarker and Its Enantiomer by Nanopore. ACS Cent. Sci. 2024, 10 (6), 1167–1178. 10.1021/acscentsci.4c00020.

(13) Kuanaeva, R. M.; Vaneev, A. N.; Gorelkin, P. V.; Erofeev, A. S. Nanopipettes as a Potential Diagnostic Tool for Selective Nanopore Detection of Biomolecules. Biosensors 2024, 14 (12), 627. 10.3390/bios14120627.

(14) Harrell, C. C.; Choi, Y.; Horne, L. P.; Baker, L. A.; Siwy, Z. S.; Martin, C. R. Resistive-Pulse DNA Detection with a Conical Nanopore Sensor. Langmuir 2006, 22 (25), 10837–10843. 10.1021/la061234k.

(15) Joty, K.; Hong, S.; Ghimire, M. L.; Kim, S.; Walker, J. N.; Brodbelt, J. S.; Yeh, H.-C.; Kim, M. J. Solid-State Nanopore Analysis of DNA-Templated Silver Nanoclusters: Voltage-Dependent Translocation and Electrolyte Stability. ACS Appl. Mater. Interfaces 2025. 10.1021/acsami.5c02405.

(16) Xue, L.; Yamazaki, H.; Ren, R.; Wanunu, M.; Ivanov, A. P.; Edel, J. B. Solid-State Nanopore Sensors. Nat Rev Mater 2020, 5 (12), 931–951. 10.1038/s41578-020-0229-6.

(17) Wu, Y.; Gooding, J. J. The Application of Single Molecule Nanopore Sensing for Quantitative Analysis. Chem. Soc. Rev. 2022, 51 (10), 3862–3885. 10.1039/D1CS00988E.

(18) Zhang, M.; Chen, C.; Zhang, Y.; Geng, J. Biological Nanopores for Sensing Applications. Proteins Structure Function and Bioinformatics 2022. 10.1002/prot.26308.

(19) Jeong, K. B.; Ryu, M.; Kim, J. S.; Kim, M.; Yoo, J.; Chung, M.; Oh, S.; Jo, G.; Lee, S. G.; Kim, H. M.; Lee, M. K.; Chi, S. W. Single-Molecule Fingerprinting of Protein-Drug Interaction Using a Funneled Biological Nanopore. Nature Communications 2023, 14 (1), 1461. 10.1038/s41467-023-37098-4.

(20) Sauciuc, A.; Morozzo della Rocca, B.; Tadema, M. J.; Chinappi, M.; Maglia, G. Translocation of Linearized Full-Length Proteins through an Engineered Nanopore under Opposing Electrophoretic Force. Nat Biotechnol 2024, 42 (8), 1275–1281. 10.1038/s41587-023-01954-x.

(21) Ahmad, M.; Ha, J.-H.; Mayse, L. A.; Presti, M. F.; Wolfe, A. J.; Moody, K. J.; Loh, S. N.; Movileanu, L. A Generalizable Nanopore Sensor for Highly Specific Protein Detection at Single-Molecule Precision. Nat Commun 2023, 14 (1), 1374. 10.1038/s41467-023-36944-9.

(22) Tang, Z.; Zhang, D.; Cui, W.; Zhang, H.; Pang, W.; Duan, X. Fabrications, Applications and Challenges of Solid-State Nanopores: A Mini Review. Nanomaterials and Nanotechnology 2016. 10.5772/64015.

(23) Storm, A. J.; Storm, C.; Chen, J.; Zandbergen, H.; Joanny, J.-F.; Dekker, C. Fast DNA Translocation through a Solid-State Nanopore. Nano Lett. 2005, 5 (7), 1193–1197. 10.1021/nl048030d.

(24) Kwok, H.; Briggs, K.; Tabard-Cossa, V. Nanopore Fabrication by Controlled Dielectric Breakdown. Plos One 2014. 10.1371/journal.pone.0092880.

(25) Fried, J. P.; Swett, J. L.; Nadappuram, B. P.; Mol, J. A.; Edel, J. B.; Ivanov, A. P.; Yates, J. R. In Situ Solid-State Nanopore Fabrication. Chem. Soc. Rev. 2021, 50 (8), 4974–4992. 10.1039/D0CS00924E.

(26) Bafna, J. A.; Soni, G. V. Fabrication of Low Noise Borosilicate Glass Nanopores for Single Molecule Sensing. PLoS ONE 2016, 11 (6), e0157399. 10.1371/journal.pone.0157399.

(27) Pal, S.; Huttner, D.; Verma, N. C.; Nemirovsky, T.; Ziv, O.; Sher, N.; Yivgi-Ohana, N.; Meller, A. Amplification-Free Quantification of Endogenous Mitochondrial DNA Copy Number Using Solid-State Nanopores. ACS Nano 2025, 19 (11), 11390–11402. 10.1021/acsnano.5c00732.

(28) Guan, X.; Li, H.; Chen, L.; Qi, G.; Jin, Y. Glass Capillary-Based Nanopores for Single Molecule/Single Cell Detection. Acs Sensors 2023. 10.1021/acssensors.2c02102.

(29) Qian, J.; Li, H.; Sun, L. The Translocation of a Polymer Through a Nanopore With Sandglass-like Geometry. Journal of Polymer Science 2023. 10.1002/pol.20230466.

(30) Yin, B.; Fang, S.; Wu, B.; Ma, W.; Zhou, D.; Yin, Y.; Tian, R.; He, S.; Huang, J.-A.; Xie, W.; Zhang, X.-H.; Wang, Z.; Wang, D. Directly Characterizing the Capture Radius of Tethered Double-Stranded DNA by Single-Molecule Nanopipette Manipulation. ACS Nano 2024, 18 (41), 27962–27973. 10.1021/acsnano.4c05605.

(31) Zhu, Z.; Zhou, Y.; Xu, X.; Wu, R.; Jin, Y.; Li, B. Adaption of a Solid-State Nanopore to Homogeneous DNA Organization Verification and Label-Free Molecular Analysis Without Covalent Modification. Analytical Chemistry 2017. 10.1021/acs.analchem.7b03442.

(32) Chen, K.; Bell, N. A. W.; Kong, J.; Tian, Y.; Keyser, U. F. Direction- and Salt-Dependent Ionic Current Signatures for DNA Sensing with Asymmetric Nanopores. Biophysical Journal 2017, 112 (4), 674–682. 10.1016/j.bpj.2016.12.033.

(33) Zheng, F.; Han, Q. Distinct DNA Conformations during Forward and Backward Translocations through a Conical Nanopore. Analyst 2024, 10.1039.D4AN01068J. 10.1039/D4AN01068J.

(34) Bell, N. A. W.; Chen, K.; Ghosal, S.; Ricci, M.; Keyser, U. F. Asymmetric Dynamics of DNA Entering and Exiting a Strongly Confining Nanopore. Nat Commun 2017, 8 (1), 380. 10.1038/s41467-017-00423-9.

(35) Zheng, F.; Alawami, M.; Zhu, J.; Platnich, C. M.; Sha, J.; Keyser, U. F.; Chen, K. DNA Carrier-Assisted Molecular Ping-Pong in an Asymmetric Nanopore. Nano Lett. 2023, 23 (23), 11145–11151. 10.1021/acs.nanolett.3c03605.

(36) Mi, Z.; Zhao, X.; Chen, X.; Wang, Y.; Shan, X.; Chen, K.; Lu, X. Ten Thousand Recaptures of a Single DNA Molecule in a Nanopore and Variance Reduction of Translocation Characteristics. ACS Nano 2024, 18 (34), 23243–23252. 10.1021/acsnano.4c05959.

(37) Plesa, C.; Cornelissen, L.; Tuijtel, M. W.; Dekker, C. Non-Equilibrium Folding of Individual DNA Molecules Recaptured up to 1000 Times in a Solid State Nanopore. Nanotechnology 2013, 24 (47), 475101. 10.1088/0957-4484/24/47/475101.

(38) Al-Waqfi, R. A.; Khan, C. J.; Irving, O. J.; Matthews, L.; Albrecht, T. Crowding Effects during DNA Translocation in Nanopipettes. ACS Nano 2025, 19 (17), 16803–16812. 10.1021/acsnano.5c01529.

(39) Harrell, C. C.; Choi, Y.; Horne, L. P.; Baker, L. A.; Siwy, Z. S.; Martin, C. R. Resistive-Pulse DNA Detection with a Conical Nanopore Sensor. Langmuir 2006, 22 (25), 10837–10843. 10.1021/la061234k.

(40) Wen, C.; Zeng, S.; Li, S.; Zhang, Z.; Zhang, S.-L. On Rectification of Ionic Current in Nanopores. Anal. Chem. 2019, 91 (22), 14597–14604. 10.1021/acs.analchem.9b03685.

(41) Deamer, D.; Akeson, M.; Branton, D. Three Decades of Nanopore Sequencing. Nat Biotechnol 2016, 34 (5), 518–524. 10.1038/nbt.3423.

(42) Kasianowicz, J. J.; Bezrukov, S. M. On “Three Decades of Nanopore Sequencing.” Nat Biotechnol 2016, 34 (5), 481–482. 10.1038/nbt.3570.

(43) Steinbock, L. J.; Bulushev, R. D.; Krishnan, S.; Raillon, C.; Radenovic, A. DNA Translocation through Low-Noise Glass Nanopores. ACS Nano 2013, 7 (12), 11255–11262. 10.1021/nn405029j.

(44) Baldelli, M.; Di Muccio, G.; Sauciuc, A.; Morozzo della Rocca, B.; Viola, F.; Balme, S.; Bonini, A.; Maglia, G.; Chinappi, M. Controlling Electroosmosis in Nanopores Without Altering the Nanopore Sensing Region. Advanced Materials 2024, 36 (33), 2401761. 10.1002/adma.202401761.

(45) Steinbock, L. J.; Steinbock, J. F.; Radenovic, A. Controllable Shrinking and Shaping of Glass Nanocapillaries under Electron Irradiation. Nano Lett. 2013, 13 (4), 1717–1723. 10.1021/nl400304y.

(46) Maheshwaram, S. K.; Sreenivasa, K.; Soni, G. V. Fingerprinting Branches on Supercoiled Plasmid DNA Using Quartz Nanocapillaries. Nanoscale 2021, 13 (1), 320–331. 10.1039/D0NR06219G.

(47) Smeets, R. M. M.; Keyser, U. F.; Krapf, D.; Wu, M.-Y.; Dekker, N. H.; Dekker, C. Salt Dependence of Ion Transport and DNA Translocation through Solid-State Nanopores. Nano Lett. 2006, 6 (1), 89–95. 10.1021/nl052107w.

(48) Conformation-Dependent Dynamics of Polymer Capture and Translocation through Solid-State Nanopores - Li - Small - Wiley Online Library. https://onlinelibrary.wiley.com/doi/10.1002/smll.202501362 (accessed 2025-07-09).

(49) Mensing, P.; Charron, M.; Bouhamidi, M. Y.; Tabard-Cossa, V. Folding Dynamics of Linear ssDNA– dsDNA Heterostructures through Solid-State Nanopores. Macromolecules 2025, 58 (8), 4094–4102. 10.1021/acs.macromol.5c00119.

(50) Mihovilovic, M.; Hagerty, N.; Stein, D. Statistics of DNA Capture by a Solid-State Nanopore. Phys. Rev. Lett. 2013, 110 (2), 028102. 10.1103/PhysRevLett.110.028102.

(51) Storm, A. J.; Chen, J. H.; Zandbergen, H. W.; Dekker, C. Translocation of Double-Strand DNA through a Silicon Oxide Nanopore. Phys. Rev. E 2005, 71 (5), 051903. 10.1103/PhysRevE.71.051903.

(52) Briggs, K.; Madejski, G.; Magill, M.; Kastritis, K.; De Haan, H. W.; McGrath, J. L.; Tabard-Cossa, V. DNA Translocations through Nanopores under Nanoscale Preconfinement. Nano Lett. 2018, 18 (2), 660–668. 10.1021/acs.nanolett.7b03987.

(53) Fraccari, R. L.; Ciccarella, P.; Bahrami, A.; Carminati, M.; Ferrari, G.; Albrecht, T. High-Speed Detection of DNA Translocation in Nanopipettes. Nanoscale 2016, 8 (14), 7604–7611. 10.1039/C5NR08634E.

(54) Albrecht, T. Single-Molecule Analysis with Solid-State Nanopores. Annual Rev. Anal. Chem. 2019, 12 (1), 371–387. 10.1146/annurev-anchem-061417-125903.

(55) Li, J.; Gershow, M.; Stein, D.; Brandin, E.; Golovchenko, J. A. DNA Molecules and Configurations in a Solid-State Nanopore Microscope. Nature Mater 2003, 2 (9), 611–615. 10.1038/nmat965.

(56) Chen, P.; Gu, J.; Brandin, E.; Kim, Y.-R.; Wang, Q.; Branton, D. Probing Single DNA Molecule Transport Using Fabricated Nanopores. Nano Lett. 2004, 4 (11), 2293–2298. 10.1021/nl048654j.

(57) Kumar Sharma, R.; Agrawal, I.; Dai, L.; Doyle, P. S.; Garaj, S. Complex DNA Knots Detected with a Nanopore Sensor. Nat Commun 2019, 10 (1), 4473. 10.1038/s41467-019-12358-4.

(58) Frykholm, K.; Müller, V.; Kk, S.; Dorfman, K. D.; Westerlund, F. DNA in Nanochannels: Theory and Applications. Quarterly Reviews of Biophysics 2022, 55, e12. 10.1017/S0033583522000117.

(59) Fologea, D.; Gershow, M.; Ledden, B.; McNabb, D. S.; Golovchenko, J. A.; Li, J. Detecting Single Stranded DNA with a Solid State Nanopore. Nano Lett 2005, 5 (10), 1905–1909. 10.1021/nl051199m.

(60) Plesa, C.; Verschueren, D.; Pud, S.; van der Torre, J.; Ruitenberg, J. W.; Witteveen, M. J.; Jonsson, M. P.; Grosberg, A. Y.; Rabin, Y.; Dekker, C. Direct Observation of DNA Knots Using a Solid-State Nanopore. Nature Nanotech 2016, 11 (12), 1093–1097. 10.1038/nnano.2016.153.

