## Supplementary File for "Reversible DNA Translocation as a molecular caliper to probe the nanoscale asymmetry of glass nanopores"

Sukanya Sadhu and G. V. Soni
Raman Research Institute, Bangalore, INDIA

SUPPLEMENTARY INFORMATION

This PDF file includes:
Tables S1-S9
Figures S1-S14

| Pore ID | Pore Dia. (nm) | Pore Resistance (MOhm) | DNA sample | Voltage (mV) |
|---|---|---|---|---|
| 1 | 16 | 28.9 | 10 kbp lin | ±300, ±500, ±700, ±900 |
| | | | 5 kbp lin | ±300, ±500, ±700, ±900 |
| | | | 3 kbp lin | ±300, ±500, ±700, ±900 |
| 2 | 18 | 33.27 | 10 kbp lin | ±300, ±500, ±700, ±900 |
| | | | 5 kbp lin | ±300, ±500, +700 |
| 3 | 19 | 15.27 | 10 kbp lin | ±300, ±500, ±700, ±900 |
| | | | 5 kbp lin | ±300, ±500, ±700, ±900 |
| | | | 3 kbp lin | ±300, ±500, ±700, ±900 |
| 4 | 17 | 14.49 | 10 kbp lin | ±300, ±500, ±700, ±900 |
| | | | 5 kbp lin | ±300, ±500, ±700, ±900 |
| | | | 3 kbp lin | ±300, ±500, ±700, ±900 |
| 5 | 19 | 20.07 | 10 kbp lin | ±300, ±500, ±700, ±900 |
| | | | 5 kbp lin | ±300, ±500, ±700, ±900 |
| 6 | 17 | 29.77 | 5 kbp lin | ±300, ±500, +700 |
| | | | 3 kbp lin | ±300, ±500, ±700, ±900 |
| 7 | 18 | 17.5 | 10 kbp lin | ±300, ±500, ±700, ±900 |
| | | | 5 kbp lin | ±300, ±500, ±700, ±900 |
| | | | 3 kbp lin | ±300, ±500, ±700, ±900 |
| 8 | 20 | 13.5 | 10 kbp lin | ±300, ±500, ±700, ±900 |
| | | | 5 kbp lin | ±300, ±500, ±700, ±900 |
| | | | 3 kbp lin | ±500, ±700, ±900 |
| 9 | 19 | 22.51 | 5 kbp lin | +300 |
| | | | 3 kbp lin | ±300, ±500, ±700, ±900 |
| 10 | 17 | 17.55 | 3 kbp lin | ±300, ±500, ±700, ±900 |

**Table S1: Data Summary.** Experimental parameters of all datasets presented in this work.

| Pore ID | DNA used | Voltage (mV) | Forward | | Reverse | | Change in $\Delta G_1$ (%) | Change in $\Delta G_2$ (%) |
|---|---|---|---|---|---|---|---|---|
| | | | $\Delta G_1$ (nS) | $\Delta G_2$ (nS) | $\Delta G_1$ (nS) | $\Delta G_2$ (nS) | | |
| 1 | 10 kbp | 300 | 0.72 ± 0.04 | 1.19 ± 0.07 | 0.68 ± 0.03 | 1.12 ± 0.05 | 5.8 | 5.4 |
| | | 500 | 0.66 ± 0.03 | 1.17 ± 0.05 | 0.59 ± 0.02 | 1.05 ± 0.03 | 10.4 | 9.7 |
| | | 700 | 0.62 ± 0.02 | 1.15 ± 0.04 | 0.55 ± 0.01 | 1.02 ± 0.03 | 12.3 | 11.5 |
| | | 900 | 0.61 ± 0.02 | 1.14 ± 0.04 | 0.52 ± 0.01 | 0.99 ± 0.02 | 13.6 | 13.1 |
| | 5 kbp | 300 | 0.74 ± 0.04 | 1.23 ± 0.07 | 0.67 ± 0.03 | 1.16 ± 0.05 | 8.7 | 5.7 |
| | | 500 | 0.68 ± 0.03 | 1.22 ± 0.05 | 0.59 ± 0.02 | 1.1 ± 0.04 | 13 | 9.7 |
| | | 700 | 0.66 ± 0.03 | 1.2 ± 0.05 | 0.56 ± 0.02 | 1.07 ± 0.03 | 15.5 | 10.8 |
| | | 900 | 0.64 ± 0.02 | 1.21 ± 0.05 | 0.53 ± 0.02 | 1.05 ± 0.03 | 17 | 13.1 |
| | 3 kbp | 300 | 0.74 ± 0.04 | 1.24 ± 0.08 | 0.67 ± 0.03 | 1.17 ± 0.06 | 9.9 | 5.3 |
| | | 500 | 0.69 ± 0.03 | 1.23 ± 0.05 | 0.59 ± 0.02 | 1.11 ± 0.04 | 14.6 | 10 |
| | | 700 | 0.66 ± 0.03 | 1.21 ± 0.05 | 0.55 ± 0.02 | 1.06 ± 0.04 | 16.7 | 12 |
| | | 900 | 0.64 ± 0.03 | 1.21 ± 0.05 | 0.53 ± 0.02 | 1.04 ± 0.04 | 17.7 | 14.1 |
| 7 | 10 kbp | 300 | 0.93 ± 0.04 | 1.62 ± 0.08 | 0.87 ± 0.03 | 1.54 ± 0.04 | 6.9 | 5.5 |
| | | 500 | 0.79 ± 0.03 | 1.49 ± 0.05 | 0.75 ± 0.02 | 1.38 ± 0.04 | 5.7 | 7.2 |
| | | 700 | 0.82 ± 0.04 | 1.54 ± 0.05 | 0.75 ± 0.02 | 1.45 ± 0.04 | 8.2 | 6.3 |
| | | 900 | 0.78 ± 0.03 | 1.5 ± 0.05 | 0.72 ± 0.02 | 1.41 ± 0.04 | 7.9 | 6.2 |
| | 5 kbp | 300 | 0.94 ± 0.04 | 1.64 ± 0.08 | 0.86 ± 0.03 | 1.52 ± 0.06 | 9.2 | 7.5 |
| | | 500 | 0.85 ± 0.03 | 1.57 ± 0.06 | 0.76 ± 0.02 | 1.43 ± 0.04 | 10.9 | 8.9 |
| | | 700 | 0.82 ± 0.04 | 1.53 ± 0.06 | 0.71 ± 0.02 | 1.38 ± 0.04 | 12.7 | 9.7 |
| | | 900 | 0.8 ± 0.04 | 1.52 ± 0.06 | 0.67 ± 0.02 | 1.33 ± 0.03 | 15.6 | 12.3 |
| | 3 kbp | 300 | 0.92 ± 0.04 | 1.59 ± 0.09 | 0.87 ± 0.04 | 1.56 ± 0.06 | 5 | 2.3 |
| | | 500 | 0.82 ± 0.04 | 1.52 ± 0.06 | 0.76 ± 0.03 | 1.43 ± 0.05 | 7.1 | 5.8 |
| | | 700 | 0.79 ± 0.04 | 1.48 ± 0.06 | 0.71 ± 0.02 | 1.37 ± 0.05 | 10.5 | 7.3 |
| | | 900 | 0.77 ± 0.03 | 1.46 ± 0.07 | 0.68 ± 0.02 | 1.34 ± 0.04 | 11.6 | 8.3 |
| 3 | 10 kbp | 300 | 0.84 ± 0.03 | 1.44 ± 0.08 | 0.8 ± 0.03 | 1.39 ± 0.05 | 4.6 | 3.9 |
| | | 500 | 0.75 ± 0.03 | 1.34 ± 0.06 | 0.69 ± 0.02 | 1.28 ± 0.04 | 8.3 | 4.4 |
| | | 700 | 0.7 ± 0.04 | 1.31 ± 0.05 | 0.65 ± 0.01 | 1.23 ± 0.03 | 7.9 | 5.9 |
| | | 900 | 0.69 ± 0.03 | 1.27 ± 0.06 | 0.62 ± 0.02 | 1.21 ± 0.03 | 9.1 | 4.9 |
| | 5 kbp | 300 | 0.83 ± 0.04 | 1.42 ± 0.07 | 0.78 ± 0.03 | 1.38 ± 0.06 | 5.7 | 2.8 |
| | | 500 | 0.73 ± 0.03 | 1.34 ± 0.05 | 0.69 ± 0.02 | 1.29 ± 0.05 | 5.3 | 3.9 |
| | | 700 | 0.69 ± 0.03 | 1.3 ± 0.05 | 0.65 ± 0.02 | 1.25 ± 0.04 | 6.3 | 3.6 |
| | | 900 | 0.68 ± 0.03 | 1.28 ± 0.05 | 0.62 ± 0.01 | 1.21 ± 0.03 | 8.1 | 5.1 |
| | 3 kbp | 300 | 0.83 ± 0.04 | 1.43 ± 0.09 | 0.78 ± 0.03 | 1.4 ± 0.06 | 6.5 | 2 |
| | | 500 | 0.76 ± 0.04 | 1.4 ± 0.07 | 0.7 ± 0.03 | 1.32 ± 0.05 | 7.4 | 5.4 |
| | | 700 | 0.73 ± 0.03 | 1.36 ± 0.08 | 0.66 ± 0.02 | 1.28 ± 0.05 | 9.6 | 5.7 |
| | | 900 | 0.71 ± 0.03 | 1.32 ± 0.08 | 0.63 ± 0.02 | 1.06 ± 0.26 | 11 | 20 |
| **Average decrease in ΔG values (%) in reverse compared to forward** | | | | | | | **9.9** | **7.6** |

**Table S2: ΔG values in forward and reverse direction.** Data is shown for Pore-1, Pore-7 & Pore-3. We can see a decrease in both $\Delta G_1$ and $\Delta G_2$ values in reverse translocation across all voltages and DNA lengths. The data has been used in Figure 3, Figure S4.

| | | C_forward | C_forward per bp | C_reverse | C_reverse per bp |
|---|---|---|---|---|---|
| P1 | 3 kbp DNA | 124.18 | 0.04 | 152.06 | 0.05 |
| | 5 kbp DNA | 179.01 | 0.04 | 277.66 | 0.05 |
| | 10 kbp DNA | 321.86 | 0.03 | 743.12 | 0.08 |
| P3 | 3 kbp DNA | 107.04 | 0.04 | 107.54 | 0.04 |
| | 5 kbp DNA | 206.11 | 0.04 | 252.75 | 0.05 |
| | 10 kbp DNA | 414.05 | 0.04 | 546.92 | 0.06 |
| P4 | 3 kbp DNA | 133.55 | 0.04 | 166.99 | 0.06 |
| | 5 kbp DNA | 227.27 | 0.04 | 317.77 | 0.06 |
| | 10 kbp DNA | 426.88 | 0.04 | 747.41 | 0.08 |

**Table S3: Coefficients of the fit of Figure 4c to equation y = C/x.** Fitting values of C_forward and C_reverse are listed for different DNA lengths measured in different nanopores. C values per base-pair are also evaluated by dividing by total number of basepairs in the DNA.

| Pore ID | Voltage (mV) | $\frac{\Delta t_r}{\Delta t_f}$ | | |
|---|---|---|---|---|
| | | **10 kbp DNA** | **5 kbp DNA** | **3 kbp DNA** |
| 1 | 300 | 2.18 | 1.51 | 1.19 |
| | 500 | 2.36 | 1.58 | 1.22 |
| | 700 | 2.32 | 1.52 | 1.19 |
| | 900 | 2.38 | 1.43 | 1.22 |
| 3 | 300 | 1.37 | 1.26 | 1.01 |
| | 500 | 1.4 | 1.2 | 1.02 |
| | 700 | 1.25 | 1.21 | 0.99 |
| | 900 | 1.22 | 1.16 | 1.03 |
| 4 | 300 | 2.03 | 1.46 | 1.28 |
| | 500 | 1.57 | 1.36 | 1.19 |
| | 700 | 1.74 | 1.38 | 1.13 |
| | 900 | 1.57 | 1.28 | 1.14 |
| **Average** | | **1.8** | **1.4** | **1.1** |

**Table S4: Ratios of translocation time in reverse to forward direction ($\frac{\Delta t_r}{\Delta t_f}$) across multiple voltages.** Data is shown for Pore-1, Pore-3 & Pore-4. The data is used in Figure 4d, Figure S6b and S6d.

| % of unfolded events | | | | | | | | | |
|---|---|---|---|---|---|---|---|---|---|
| | | Applied voltage | | | | | | | |
| **Pore ID** | **DNA used** | **+300 mV** | **-300 mV** | **+500 mV** | **-500 mV** | **+700 mV** | **-700 mV** | **+900 mV** | **-900 mV** |
| 1 | 10 kbp lin | 18.68 | 44.03 | 21.58 | 50.70 | 23.03 | 53.02 | 27.82 | 56.74 |
| | 5 kbp lin | 24.44 | 42.71 | 30.51 | 50.82 | 32.01 | 55.41 | 37.20 | 56.64 |
| | 3 kbp lin | 33.75 | 52.38 | 37.69 | 63.17 | 45.37 | 64.40 | 47.94 | 67.86 |
| 2 | 10 kbp lin | 15.99 | 42.64 | 18.70 | 48.52 | 24.03 | 51.92 | 26.64 | 55.94 |
| | 5 kbp lin | 23.62 | 56.74 | 30.54 | -- | 37.21 | -- | -- | -- |
| 3 | 10 kbp lin | 15.92 | 38.38 | 20.78 | 42.74 | 19.91 | 46.05 | 25.29 | 36.77 |
| | 5 kbp lin | 23.48 | 40.54 | 28.01 | 43.45 | 33.73 | 49.77 | 37.34 | 49.27 |
| | 3 kbp lin | 37.18 | 51.16 | 45.75 | 56.24 | 49.28 | 59.99 | 51.33 | 57.84 |
| 4 | 10 kbp lin | 18.45 | 33.57 | 17.13 | 40.93 | 22.06 | 36.08 | 27.71 | 53.55 |
| | 5 kbp lin | 23.76 | 40.86 | 28.07 | 47.17 | 32.79 | 49.40 | 35.31 | 52.26 |
| | 3 kbp lin | 32.81 | 47.87 | 39.47 | 58.87 | 43.51 | 60.99 | 49.74 | 60.32 |
| 5 | 10 kbp lin | 16.35 | 30.99 | 23.09 | 36.74 | 24.32 | 43.48 | 26.61 | 47.09 |
| | 5 kbp lin | 20.25 | 40.69 | 30.22 | 45.74 | 33.84 | 52.67 | 40.34 | 54.27 |
| 6 | 5 kbp lin | 24.75 | 35.55 | 30.57 | -- | 34.51 | -- | -- | -- |
| | 3 kbp lin | 38.22 | 55.31 | 41.46 | 54.85 | 51.07 | 62.76 | 48.63 | 63.15 |
| 7 | 10 kbp lin | 16.17 | 25.80 | 14.58 | 35.32 | 22.88 | 47.70 | 20.09 | 49.42 |
| | 5 kbp lin | 23.32 | 37.41 | 28.85 | 45.39 | 33.50 | 51.22 | 38.98 | 54.20 |
| | 3 kbp lin | 29.60 | 45.46 | 41.52 | 50.25 | 43.75 | 57.83 | 49.28 | 59.31 |
| 8 | 10 kbp lin | 13.73 | 27.08 | 18.57 | 38.30 | 26.25 | 44.72 | 25.34 | 44.07 |
| | 5 kbp lin | 27.41 | 34.36 | 28.97 | 41.77 | 33.50 | 44.77 | 37.37 | 47.68 |
| | 3 kbp lin | -- | -- | 45.91 | 57.11 | 49.93 | 60.00 | 51.75 | 65.49 |
| 9 | 3 kbp lin | 38.67 | 52.08 | 38.24 | 58.90 | 43.09 | 62.31 | 47.68 | 65.46 |
| 10 | 3 kbp lin | 38.08 | 48.47 | 41.67 | 53.23 | 42.52 | 58.27 | 50.91 | 60.89 |

**Table S5: Percentage of unfolded events.** The total number of events is ~1000 for each experiment. This data is used in Figure 5 & Figure S8. Data shows a clear trend of the unfolded percentage increasing with applied voltage and decreasing with DNA length. It also shows excellent reproducibility across multiple pores.

**a**

| % of unfolded events | | | | |
|---|---|---|---|---|
| **DNA used** | **Applied voltage** | | | |
| | **+300 mV** | **+500 mV** | **+700 mV** | **+900 mV** |
| 10 kbp | 16.47 ± 1.56 | 19.20 ± 2.66 | 23.21 ± 1.83 | 25.64 ± 2.45 |
| 5 kbp | 23.88 ± 1.85 | 29.47 ± 1.04 | 33.89 ± 1.44 | 37.76 ± 1.57 |
| 3 kbp | 35.47 ± 3.21 | 40.83 ± 2.89 | 45.51 ± 3.24 | 49.36 ± 1.45 |

**b**

| % of unfolded events | | | | |
|---|---|---|---|---|
| **DNA used** | **Applied voltage** | | | |
| | **-300 mV** | **-500 mV** | **-700 mV** | **-900 mV** |
| 10 kbp | 34.64 ± 6.72 | 41.89 ± 5.42 | 46.14 ± 5.26 | 49.08 ± 6.62 |
| 5 kbp | 41.11 ± 6.5 | 45.72 ± 2.86 | 50.54 ± 3.26 | 52.39 ± 3.08 |
| 3 kbp | 50.39 ± 3.07 | 56.5 ± 3.69 | 60.94 ± 2.11 | 62.12 ± 3.28 |

**Table S6: Percentage of unfolded events averaged over all the nanopores used in this study.** (a) Average percentages of unfolded events across multiple pores in forward and (b) in reverse directions. The error bars present standard deviation of multiple datasets measured in multiple nanopores.

| Pore ID | Voltage (mV) | 10 kbp DNA | 5 kbp DNA | 3 kbp DNA |
|---|---|---|---|---|
| 1 | 300 | 1.84 | 1.25 | 1.01 |
| | 500 | 1.98 | 1.30 | 1.01 |
| | 700 | 2.00 | 1.28 | 1.02 |
| | 900 | 2.01 | 1.24 | 1.01 |
| 3 | 300 | 1.18 | 1.13 | 0.93 |
| | 500 | 1.12 | 1.09 | 0.97 |
| | 700 | 1.14 | 1.07 | 0.92 |
| | 900 | 1.09 | 1.08 | 0.93 |
| 4 | 300 | 1.57 | 1.28 | 1.15 |
| | 500 | 1.44 | 1.26 | 1.12 |
| | 700 | 1.45 | 1.22 | 1.09 |
| | 900 | 1.41 | 1.24 | 1.13 |

**Table S7: ECD ratios ($\frac{ECD_r}{ECD_f}$) of different DNA at different voltages.** ECD ratios follow a similar trend to Δt ratios. Data is shown for Pore-1, Pore-3 & Pore-4.

| $L_{eff}$ Values (nm) | | | | | | | | | |
|---|---|---|---|---|---|---|---|---|---|
| | | **Applied voltage (mV)** | | | | | | | |
| **Pore ID** | **DNA used** | **300** | **-300** | **500** | **-500** | **700** | **-700** | **900** | **-900** |
| 1 | 10 kbp | 113.7 | 71.5 | 121.2 | 63.1 | 176.9 | 75.8 | 180.2 | 95.1 |
| 2 | | 60.3 | 38.7 | 91.5 | 67.5 | 125.4 | 89.7 | 164.7 | 128.7 |
| 3 | | 54.3 | 60.2 | 99.6 | 96.7 | 117.8 | 101.3 | 141.9 | 110 |
| 4 | | 94.5 | 60.7 | 119.4 | 99.1 | 129.2 | 85.7 | 170.6 | 146.8 |
| 5 | | 62.5 | 50.9 | 93.1 | 67.9 | 126 | 87.6 | 174.8 | 109.1 |
| 7 | | 79.1 | 57.8 | 116 | 112.3 | 138.8 | 112.4 | 132.3 | 116.6 |
| 8 | | 123.1 | 61.6 | 126.5 | 67.8 | 132.8 | 129.4 | 165.2 | 127.3 |
| **Avg ± SD** | | **83.9 ± 25.2** | **57.3 ± 9.5** | **109.6 ± 13.4** | **82.1 ± 18.5** | **135.3 ± 18.0** | **97.4 ± 17.0** | **161.4 ± 16.3** | **119.1 ± 15.5** |

**Table S8: Sensing Length ($L_{eff}$) values with voltage and translocation direction for multiple nanopores.** This data is used in Figure 7 and Figure S11-13.

| | **Result from model** | **Result from experiments** | | |
|---|---|---|---|---|
| | | **Pore 1<br>- 300 mV** | **Pore 7<br>+300 mV** | **Pore 4<br>+300 mV** |
| Slope1 | $(4.1 \pm 0.15) \times 10^{-3}$ | $(5.3 \pm 0.82) \times 10^{-3}$ | $(7.6 \pm 1.41) \times 10^{-3}$ | $(5.4 \pm 1.12) \times 10^{-3}$ |
| Slope2 | $(1.35 \pm 0.00) \times 10^{-6}$ | $(0.90 \pm 1.14) \times 10^{-5}$ | $(5.19 \pm 2.67) \times 10^{-5}$ | $(3.76 \pm 2.01) \times 10^{-5}$ |
| **$L_{eff}$** | **71.7** | **71.5** | **79.1** | **94.5** |

**Table S9: Comparison of Sensing Length ($L_{eff}$) values from experimental estimation and analytical model.**

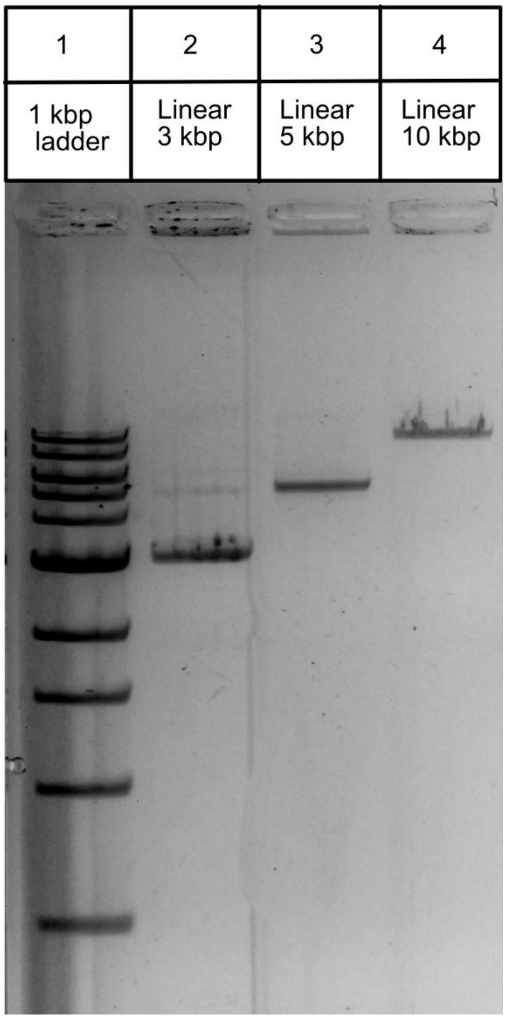


**Figure S1: Gel Electrophoresis image of 3, 5 & 10 kbp linear DNA.** Lane 1 shows a 1kb ladder. Lane 2-4 shows 3, 5 & 10 kbp linear DNA. The exact base pair numbers are 3025, 5094 and 9546 bp respectively.

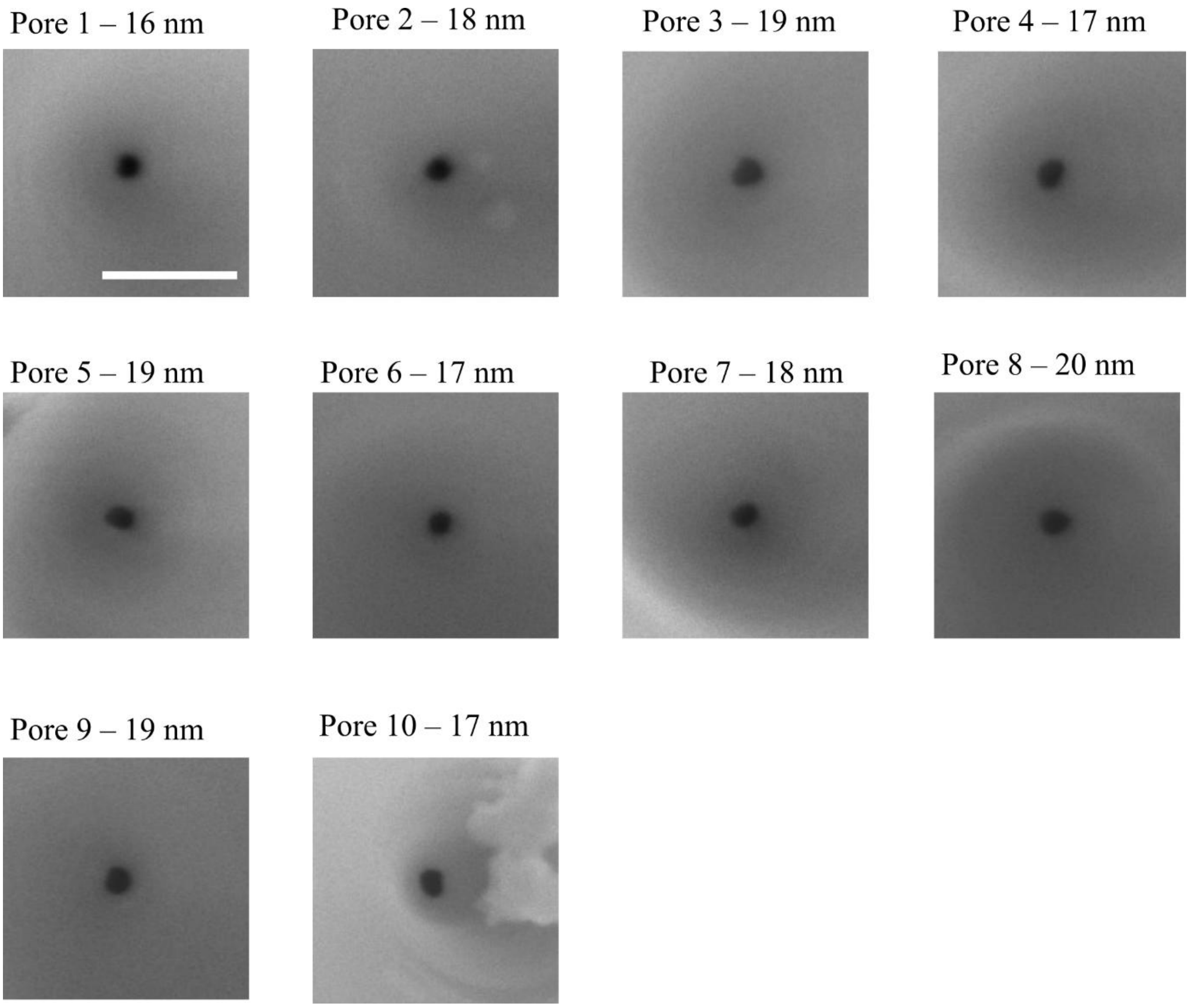


**Figure S2: SEM image of nanopores used in the experiments.** Shows the SEM images of all the nanopores used in this study. Scalebar (100 nm) is same for all images.

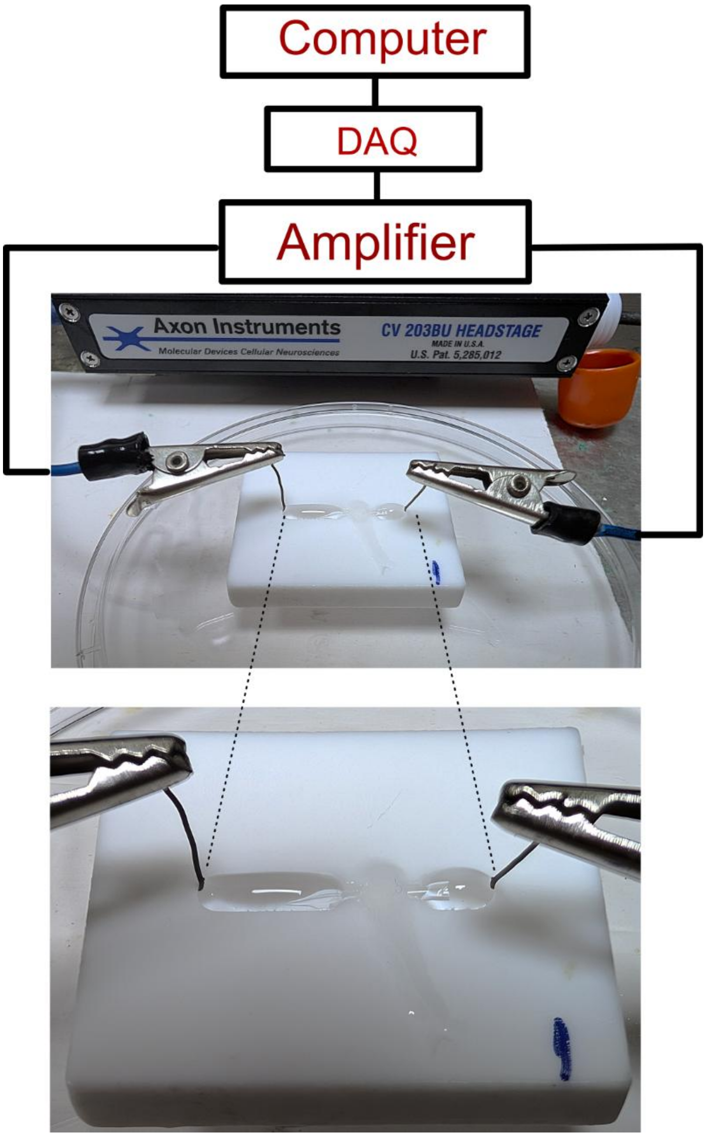


**Figure S3: Image of a typical nanopore device used in this study.** The nanopore mounted on a Teflon flow-cell (see zoomed image below) using glue. The flow cell has approximately 100 µl in volume, combining both chambers, which are filled with nanopore buffer (NPB). Two Ag/AgCl electrodes are connected across the nanopore. The DNA is added to the cis side at first for forward translocation. The voltage and current measurements are done by the Axon 200B amplifier and recorded in the PC by custom LabView software and proceeded for further analysis.

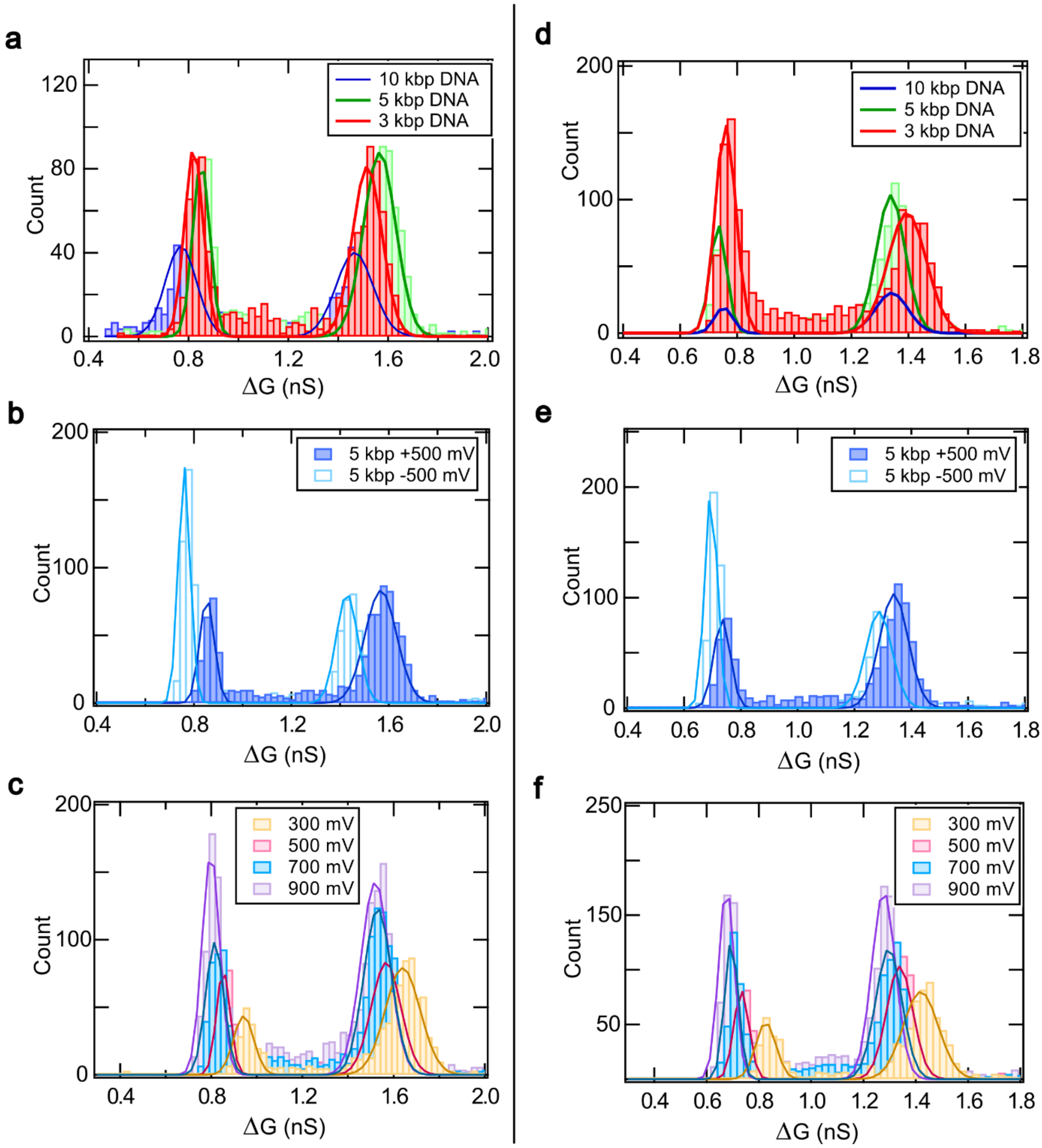


**Figure S4: Additional dataset showing reproducibility of Figure 3.** Figure a, b & c correspond to Pore-3 and Figure d, e, & f correspond to Pore-4. Plots show the Gaussian fit of ΔG data with DNA length (a & d), translocation direction (b & e) and applied voltage bias (c & f) respectively for two additional nanopores.

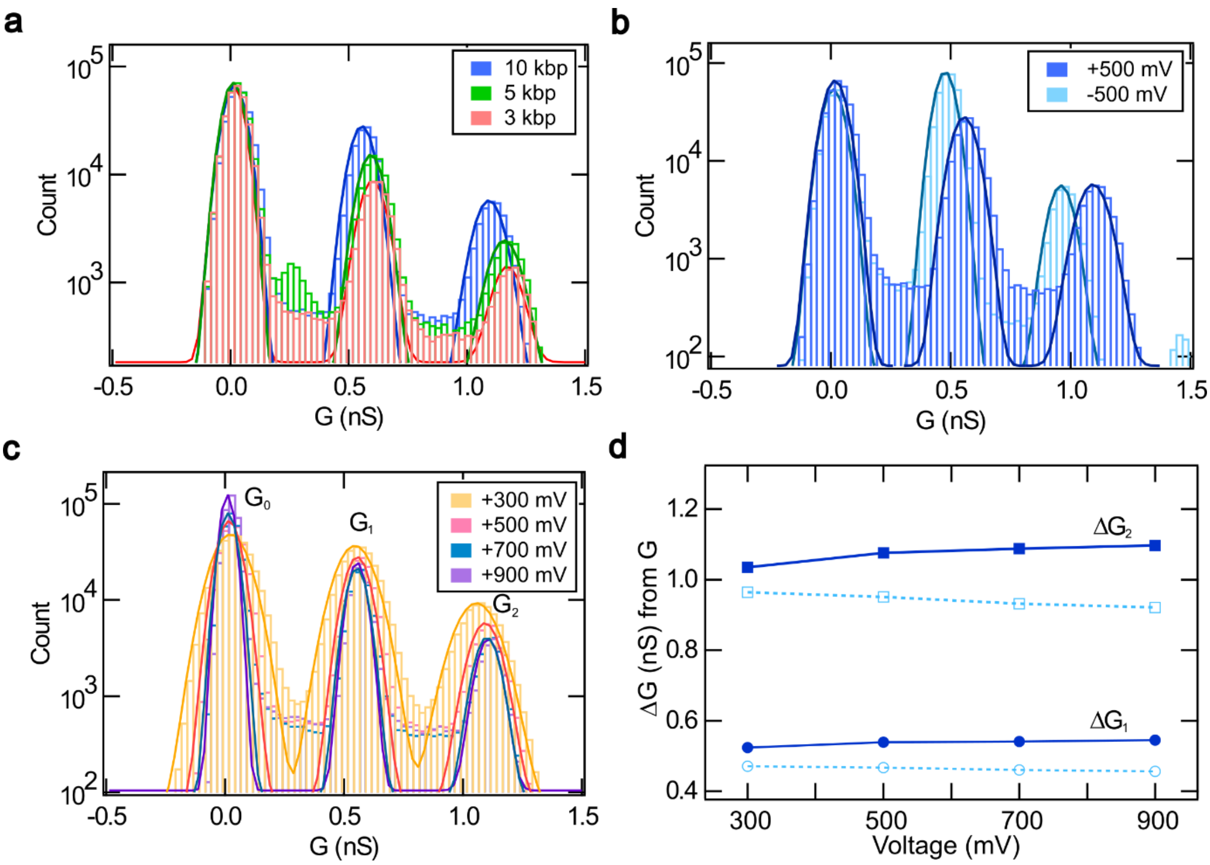


**Figure S5: Characterization of ΔG by G-histograms.** Here the ΔG values are estimated using G-histograms. (a) G-histogram for 3, 5 & 10 kbp linear DNA taken at +500 mV. The solid lines are Gaussian peak fits. (b) G-histogram for 5 kbp linear DNA at +500 mV and -500 mV. (c) G-histogram for 5 kbp linear DNA at +300 to +900 mV. (d) ΔG values calculated from the mean values of the G-histogram, $\Delta G_1 = G_1 - G_0$ & $\Delta G_2 = G_2 - G_0$, respectively. The calculated ΔG values are plotted against the voltage.

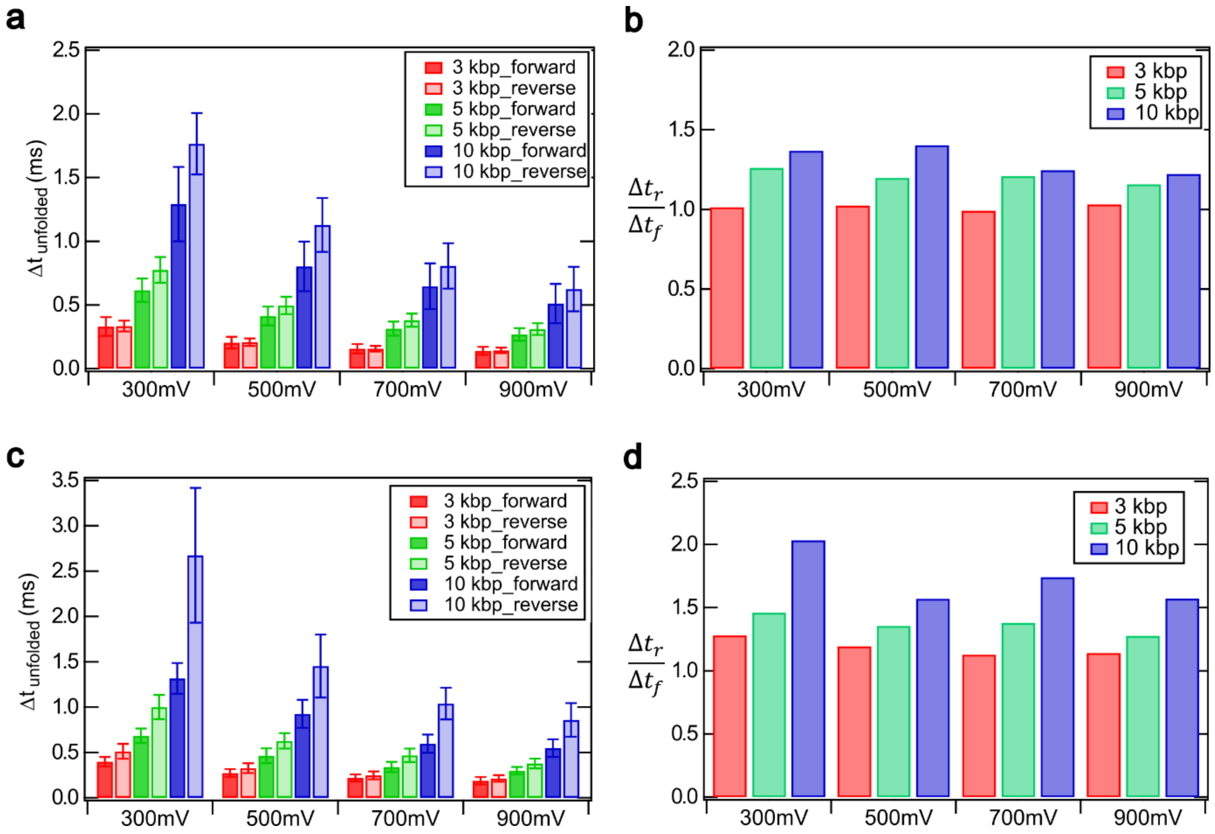


**Figure S6: Characteristics of the translocation time.** This figure shows the reproducibility of Figure 4b and 4d in Pore-3 and Pore-4. (a) & (c) Bar plots showing the mean translocation time of the unfolded population with different DNA lengths and voltages. (b) & (d) Ratio of the translocation time in the reverse to the forward direction with applied voltage and DNA length.

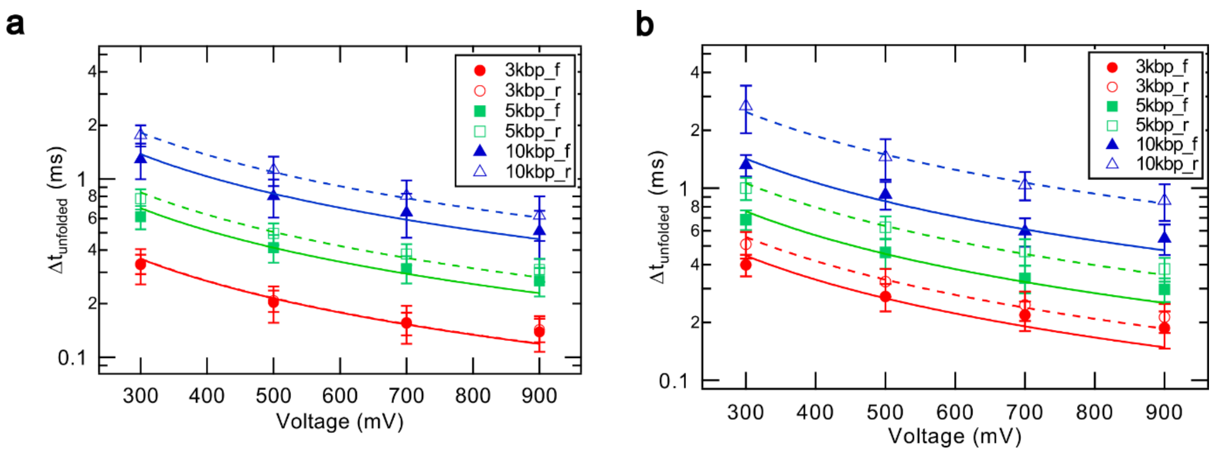


**Figure S7: Reproducibility of Figure 4c in Pore-3 and Pore-4.** (a) & (b) $\Delta t_{unfolded}$ as a function of applied voltage for 3, 5 & 10 kbp linear DNA. The data is fitted with an inverse function y~1/x and the coefficients are given in Table S3.

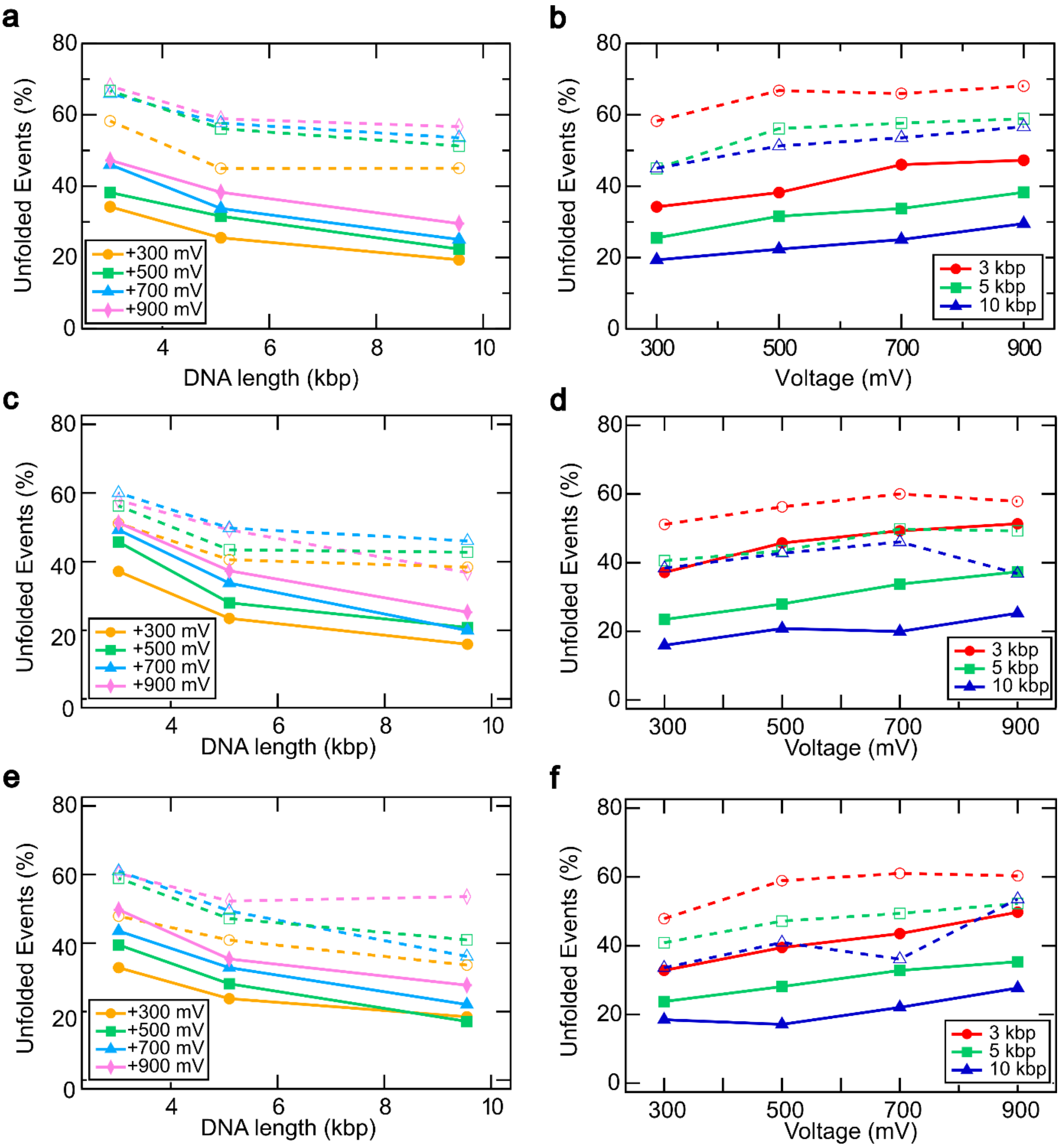


**Figure S8: Percentage of unfolded events.** (a), (c) & (e) show unfolded events decrease with DNA length. (b), (d) & (f) show unfolded event percentage increases with applied voltage. The plots show individual data of Figure 5c & 5d. The data is shown top to bottom for Pores 1, 3 & 4, respectively.

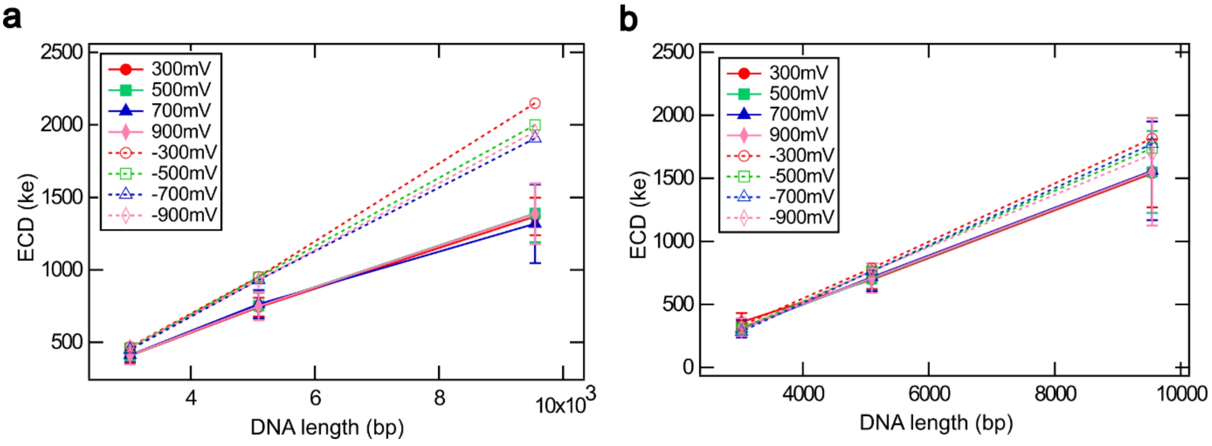


**Figure S9: ECD characteristics with DNA length.** (a) & (b) The mean ECD values plotted against 3 different DNA lengths for forward and reverse translocation.

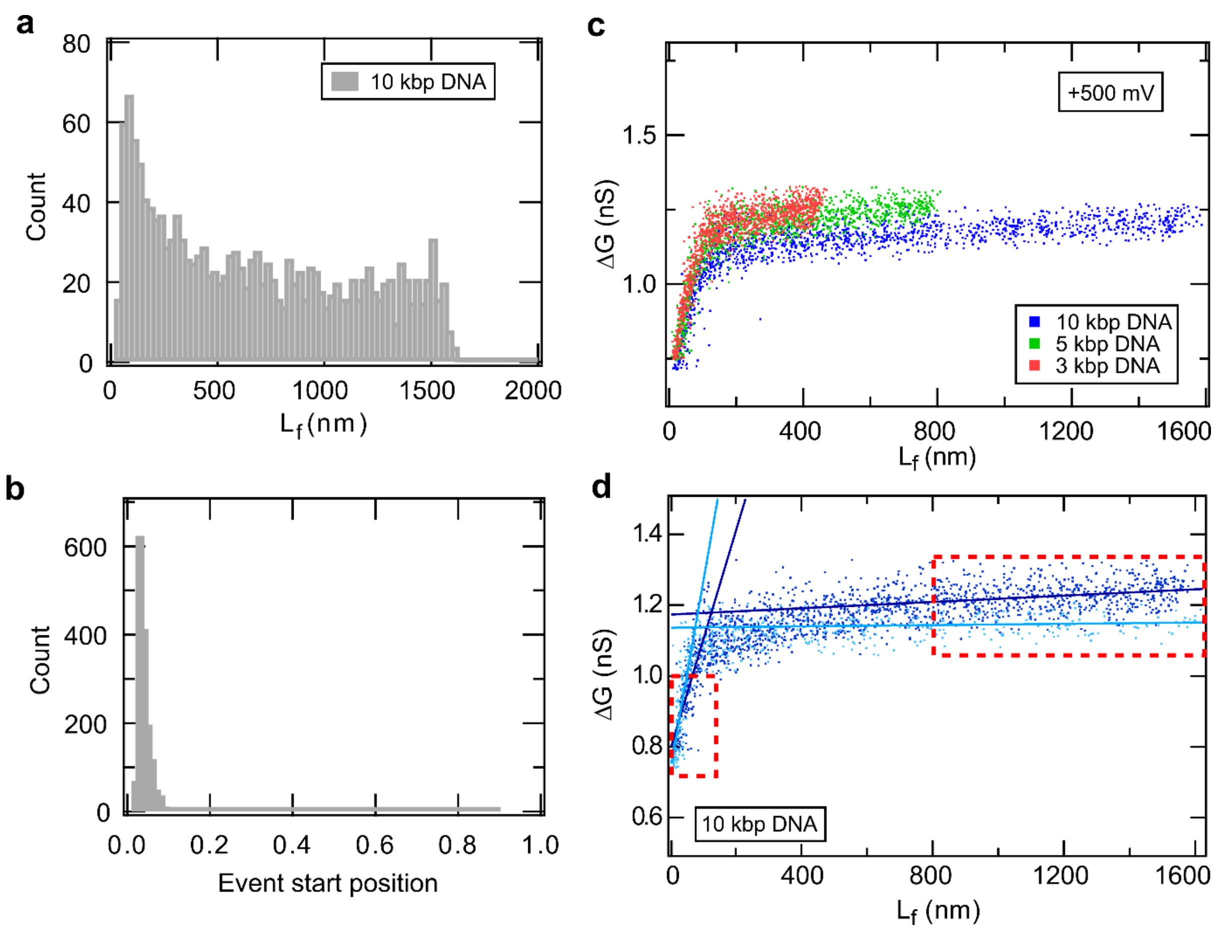


**Figure S10: Estimation of $L_{eff}$.** (a) shows histogram of DNA fold length ($L_f$) measured for 10 kbp DNA. (b) shows histogram of start position of folded region relative to the beginning of the event for 10 kbp DNA. (c) Scatter plot of ΔG vs $L_f$ for 3, 5 & 10 kbp DNA. (d) Piecewise linear fit for determination of sensing length for the 10 kbp sample in Pore-1 for +300 mV (dark blue) & -300 mV (light blue).

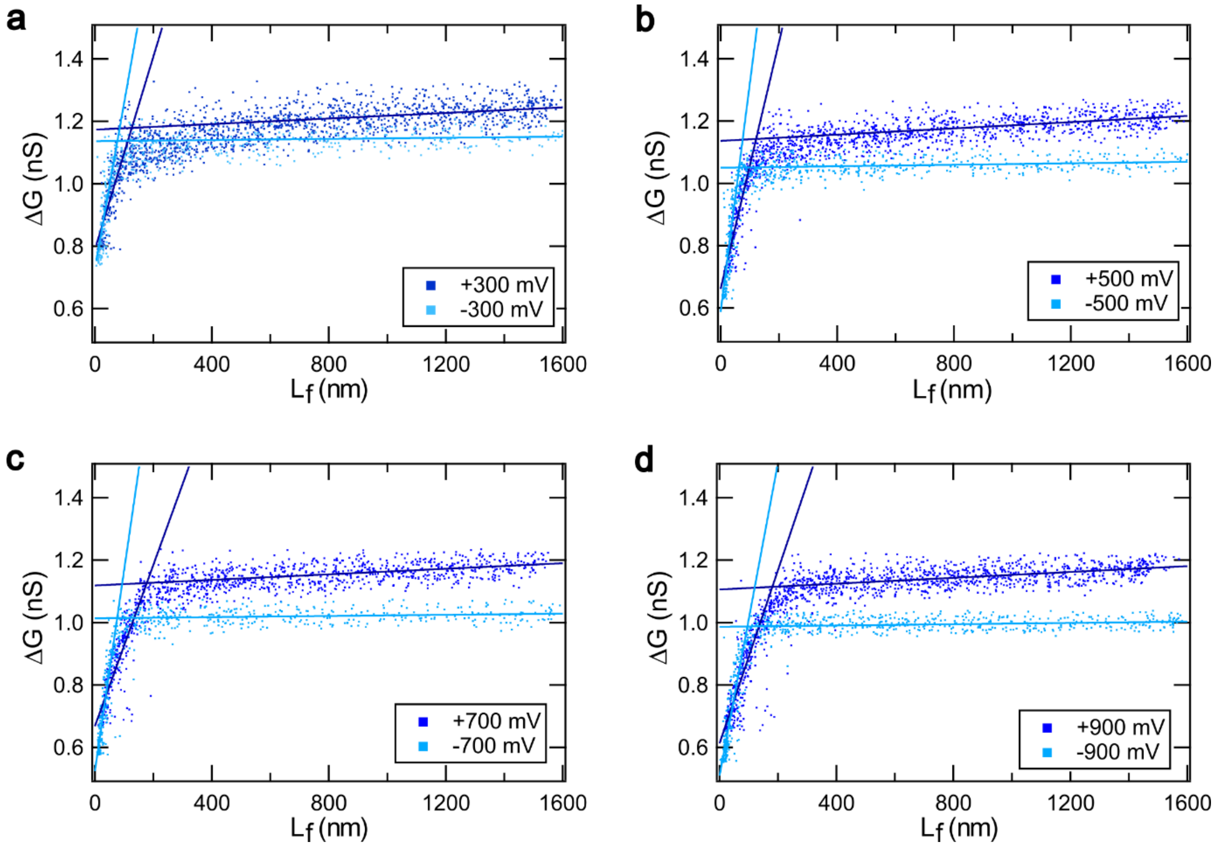


**Figure S11: ΔG vs $L_f$ plots with piecewise linear fit for multiple voltages.** Data shown for Pore-1 with 4 voltages in both forward and reverse directions (±300 mV, ±500 mV, ±700 mV, ±900 mV). The $L_{eff}$ values calculated from the fittings are given in Table S8.

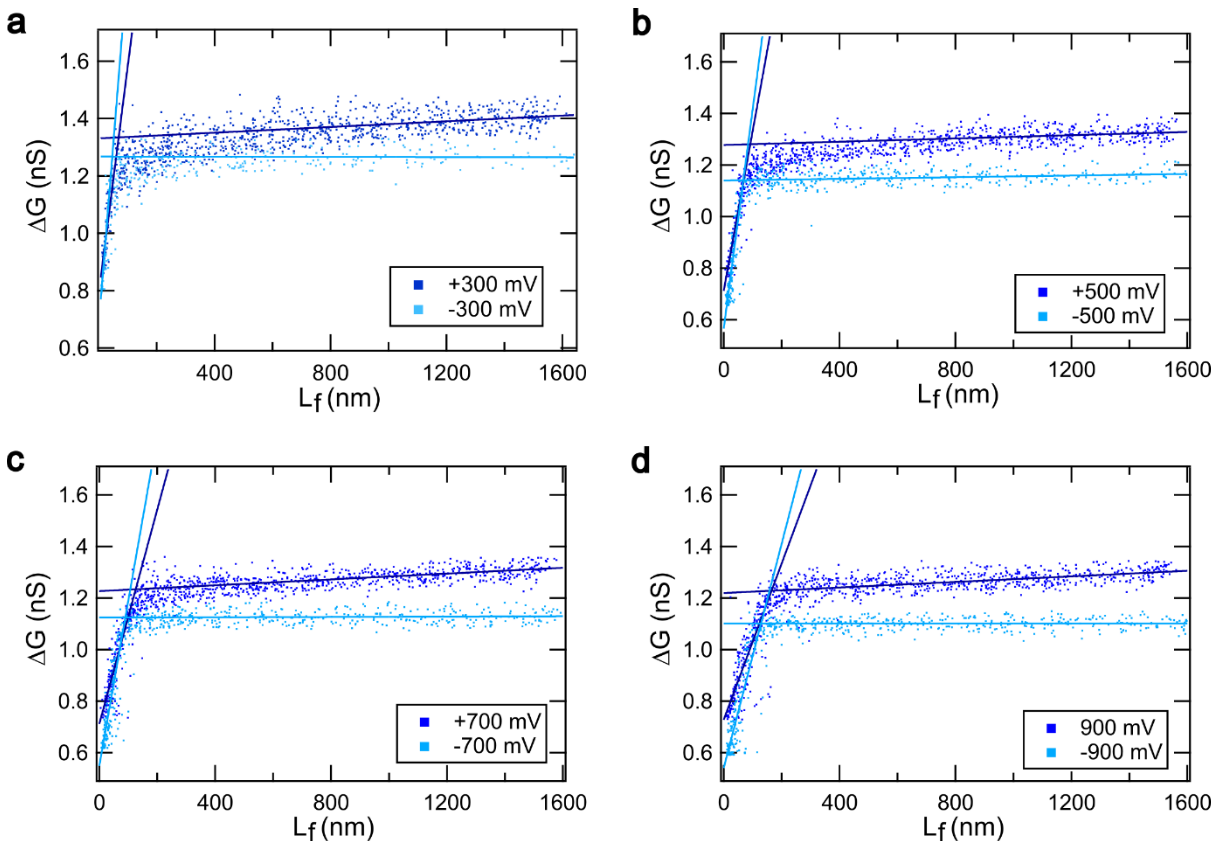


**Figure S12: ΔG vs $L_f$ plots with piecewise linear fit for multiple voltages.** Data shown for Pore-2 with 4 voltages in both forward and reverse. The $L_{eff}$ values calculated from the fittings are given in Table S8.

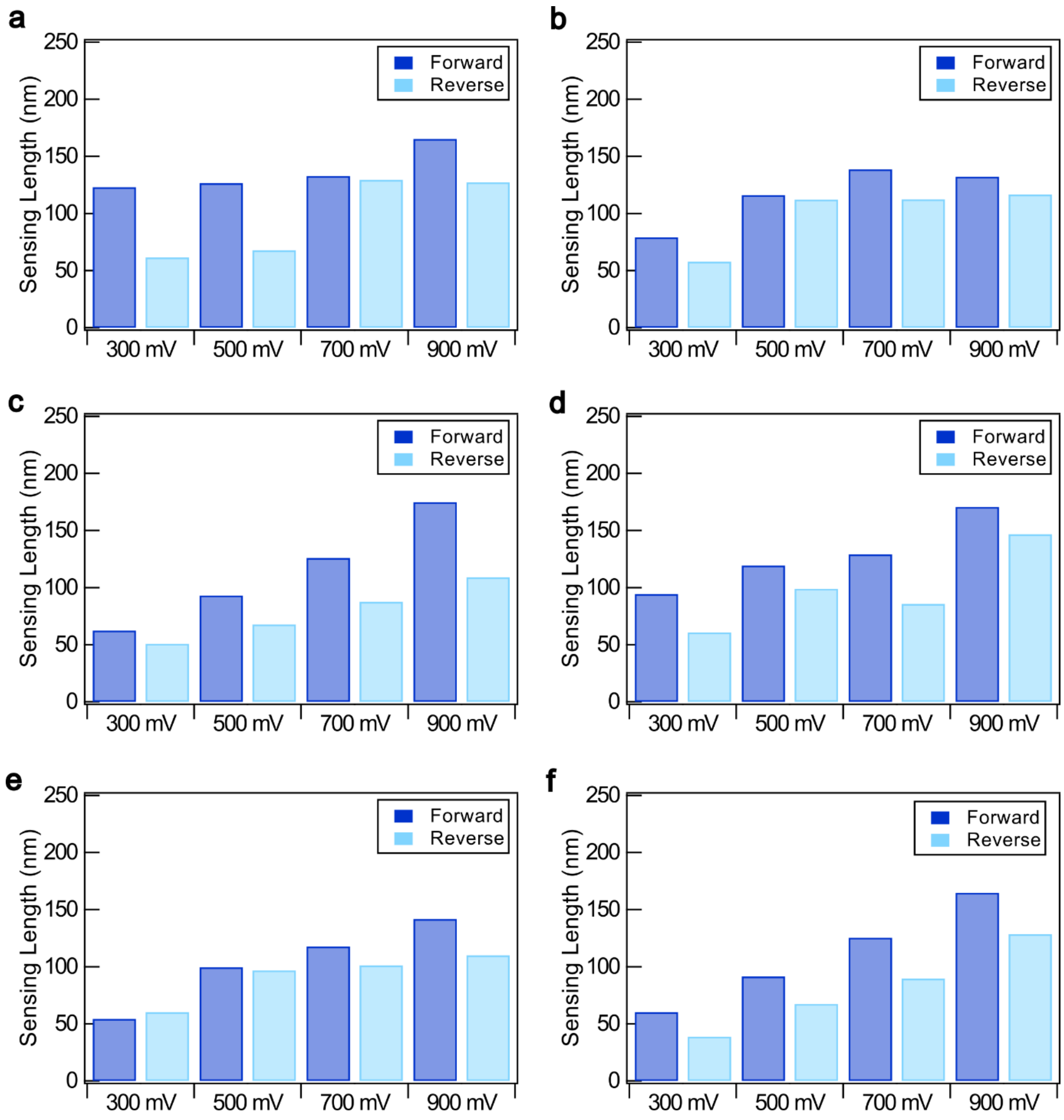


**Figure S13: Bar plots of sensing length ($L_{eff}$) values with increasing voltages across multiple nanopores.** (a)-(f) The datasets correspond to Pore 2, 3, 4, 5, 7 & 8 respectively. The values are given in Table S8.

### Details of the model for the simulation of sensing length

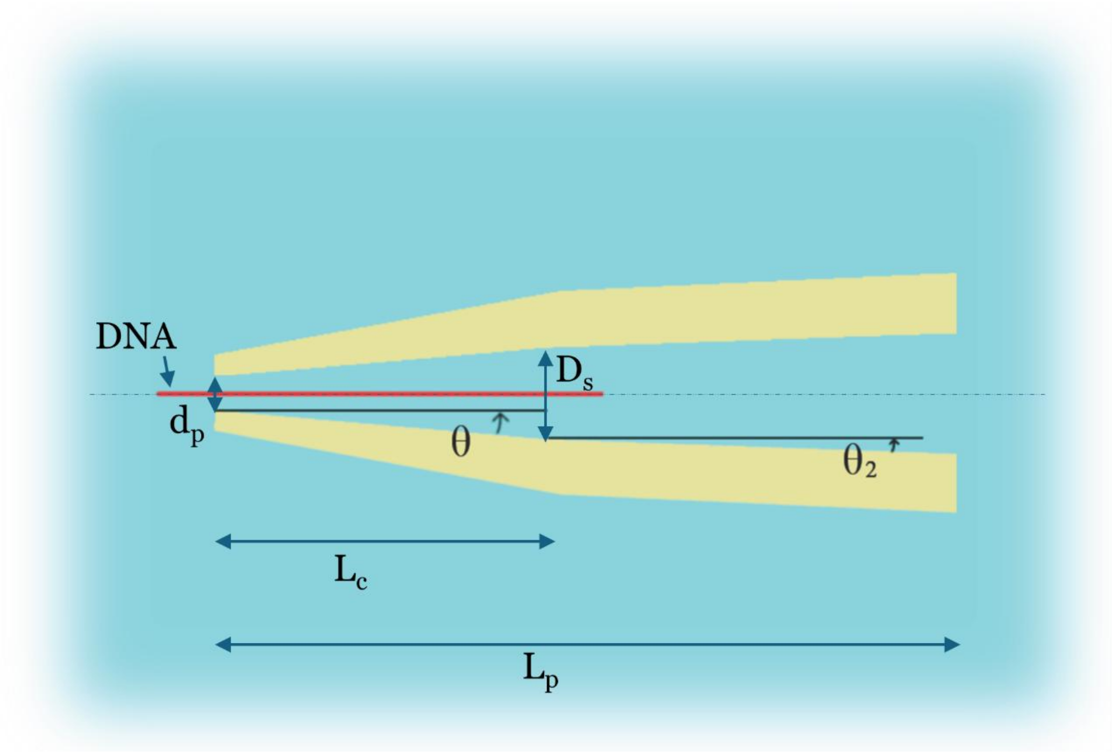


**Figure S14: Detailed schematic of the double cone model used for modelling the sensing length.** The capillary consists of two cones of angles $\theta$ and $\theta_2$. $L_C$ is the length of the first cone and $L_P$ is the pore length. The tip diameter of the nanopore is $d_P$ and the diameter at the base of the first cone is $D_s$. The DNA is shown as a cylindrical rod of diameter $d_D$. The length of the DNA is given by $L_D$.

### Details of the model for the simulation of sensing length

We have used an axially symmetric double conical model as shown in Figure S13. To simulate the sensing length of a pore ($L_{eff}$), we have analytically estimated $\Delta G$ with varying lengths of folded DNA ($L_f$). In our model, the nanopore is simplified as a connected double conical geometry with cone angles $\theta$ and $\theta_2$. The diameter of the pore at the tip is denoted by $d_P$ and the diameter of the base of the first cone is $D_s$. The DNA is considered as a solid cylinder (shown in red) with cross-sectional diameter $d_D$. $L_C$ and $L_P$ are the length of the first cone and the pore length respectively. $L_D$ is the total contour length of the DNA. $\sigma$ is the conductivity of the buffer. The values of the parameters used in our model are given below:

$\sigma$=18.05 S $m^{-1}$

$d_D$=2.5 nm

$d_P$=15 nm

$L_P$=2000 nm

$L_D$=0:1500 nm

$L_C$=300 nm

θ = 5*pi/180 = 0.087 rad

$\theta_2$ = 2*pi/180 = 0.035 rad

The **open-pore conductance ($G_P$)** of the two cones is estimated as the summation of the conductance of the individual cones, given by,

$$\frac{1}{G_p} = \frac{1}{G_{p1}} + \frac{1}{G_{p2}}$$

$$\frac{1}{G_{p1}} = R_{p1} = \sum_{i=1}^{L_c} \frac{4}{\pi\sigma}\left(\frac{1}{d_p(i)*d_p(i+1)}\right), \qquad \text{where } d_p(i) = d_P + 2i\tan\theta,$$

$$\frac{1}{G_{p2}} = R_{p2} = \sum_{i=L_c+1}^{L_P} \frac{4}{\pi\sigma}\left(\frac{1}{d_p(i)*d_p(i+1)}\right), \qquad \text{where } d_p(i) = D_s + 2i\tan\theta 2,$$

$$G_p = \frac{1}{R_{p1} + R_{p2}}$$

The **conductance with DNA ($\boldsymbol{G_{p\_WD}}$)** is given by,

$$d_p(i) = d_P + 2i\tan\theta\,, upto\ i = L_c$$

$$and, d_p(i) = d_P + 2i\tan\theta_2\,(i)\,, in\ range\ of\ i = L_c + 1\ to\ L_P$$

Case-I:

When the DNA length is shorter than the length of the first cone, $L_D < L_p$ & $L_D < L_c$,

$$\frac{1}{G_{p\_wD}} = R_{p\_wD} = \frac{4}{\sigma\pi}\sum_{i=1}^{L_D}\left(\frac{1}{\sqrt{(d_p^2(i)-2d_D^2)}*\sqrt{(d_p^2(i+1)-2d_D^2)}}\right) + \frac{4}{\sigma\pi}\sum_{i=L_D+1}^{L_c}\left(\frac{1}{\sqrt{(d_p^2(i)-d_D^2)}*\sqrt{(d_p^2(i+1)-d_D^2)}}\right) + \frac{4}{\sigma\pi}\sum_{i=L_c+1}^{L_P}\left(\frac{1}{\sqrt{(d_p^2(i)-d_D^2)}*\sqrt{(d_p^2(i+1)-d_D^2)}}\right)$$

Case-II:

When the DNA length is longer than the length of the first cone, $L_D < L_p$ & $L_D > L_c$,

$$\frac{1}{G_{p\_wD}} = R_{p\_wD} = \frac{4}{\sigma\pi}\sum_{i=1}^{L_C}\left(\frac{1}{\sqrt{(d_p^2(i)-2d_D^2)}*\sqrt{(d_p^2(i+1)-2d_D^2)}}\right) + \frac{4}{\sigma\pi}\sum_{i=L_C+1}^{L_D}\left(\frac{1}{\sqrt{(d_p^2(i)-2*d_D^2)}*\sqrt{(d_p^2(i+1)-2*d_D^2)}}\right) + \frac{4}{\sigma\pi}\sum_{i=L_D+1}^{L_P}\left(\frac{1}{\sqrt{(d_p^2(i)-d_D^2)}*\sqrt{(d_p^2(i+1)-d_D^2)}}\right)$$

Therefore, the drop in conductance (ΔG) is,

$$\Delta G = G_p - G_{p\_wD}$$

We plot the simulated ΔG with $L_f$ (Figure 7d) and fit the data using a piecewise linear function. From the intersection, we calculate the sensing length $L_{eff}$ as given in Table S7 & S8.
